# Convergent evolution of distinct immune sensor systems for fungal polygalacturonases in *Brassicaceae*

**DOI:** 10.1101/2020.12.10.418392

**Authors:** Lisha Zhang, Chenlei Hua, Rory N. Pruitt, Si Qin, Lei Wang, Isabell Albert, Markus Albert, Jan A. L. van Kan, Thorsten Nürnberger

## Abstract

Plant pattern recognition receptors (PRRs) facilitate recognition of microbial surface patterns and mediate activation of plant immunity. *Arabidopsis thaliana* RLP42, a leucine-rich repeat (LRR) receptor protein (LRR-RP), senses fungal endopolygalacturonases (PGs) through a ternary complex comprising RLP42, the adapter kinase SOBIR1, and SERK proteins. Several fungal PGs harbor a conserved 9-amino acid fragment pg9(At), which is sufficient to activate RLP42-dependent plant immunity. Domain swap experiments using RLP42 and paralogous RLP40 sequences revealed a dominant role of the island domain (ID) for ligand binding and PRR complex assembly. Involvement of the ID in plant receptor function is reminiscent of plant phytosulfokine (PSK) perception through the receptor, PSKR, a LRR receptor kinase. Sensitivity to pg9(At), which is restricted to *A. thaliana*, exhibits notable accession specificity as active RLP42 alleles were found in only 16 of 52 accessions tested. *Arabidopsis arenosa* and *Brassica rapa*, two *Brassicaceae* species closely related to *A. thaliana*, perceive plant immunogenic PG fragments pg20(Aa) or pg36(Bra), which are distinct from pg9(At). Our study unveils unprecedented complexity and dynamics of PG pattern recognition receptor evolution within a single plant family. PG perception systems may have evolved rather independently as a result of convergent evolution even among closely related species.

## Introduction

Plants employ immune receptors to detect invasive microbes and activate immunity. Detection of pathogen-associated or microbe-associated molecular patterns (PAMPs/MAMPs) by cell surface localized pattern recognition receptors (PRRs) initiates pattern-triggered immunity (PTI). This form of immunity controls attempted infection by host non-adapted microbes and contributes to basal immunity to host-adapted phytopathogens^1–3^. Host-adapted microbial invaders produce effectors to suppress PTI and to parasitize host plants. Effective immunity to these pathogens relies on effector recognition through intracellular immune receptors mediating activation of effector-triggered immunity (ETI)^4^. Defense response outputs associated with PTI or ETI largely overlap. However, programmed host cell death is a hallmark of ETI, but much less frequently observed upon PTI activation^4,5^. Mechanistic links between these two types of plant immunity have been proposed. An *A. thaliana* plasma membrane-associated intracellular hub made of helper NLRs from the ADR1 (ACTIVATED DISEASE RESISTANCE 1) family, and lipase-like proteins EDS1 (ENHANCED DISEASE SUSCEPTIBILITY 1) and PAD4 (PHYTOALEXIN DEFICIENT 4), is shared by signaling pathways activated through surface PRRs and intracellular immune receptors, and may constitute a potential link between PTI and ETI^6^.

Plant PRRs with leucine-rich repeat (LRR) ectodomains predominantly mediate recognition of proteinaceous microbe-derived patterns^1–3^. LRR-receptor kinases (LRR-RKs) contain a ligand-binding ectodomain, a single-pass transmembrane domain, and a cytoplasmic kinase domain. LRR-receptor proteins (LRR-RPs) share the same basic structure, but lack an intracellular kinase domain. LRR-RPs usually associate with the adapter LRR-RK SUPPRESSOR OF BRASSINOSTEROID INSENSITIVE 1 (BRI1)-ASSOCIATED KINASE (BAK1)-INTERACTING RECEPTOR KINASE 1 (SOBIR1) in a ligand-independent manner. The bipartite complex is structurally analogous to an LRR-RK. Upon ligand perception, both LRR-RKs and LRR-RPs recruit members of the SOMATIC EMBRYOGENESIS RECEPTOR KINASE (SERK) family of proteins as co-receptors to initiate intracellular signal transduction^1–3^. Bacterial flagellin-induced complex formation of LRR-RK FLAGELLING SENSING 2 with SERK3/BAK1 has been resolved at atomic level^7^. In contrast, mechanistic insight into ligand-induced formation of ternary complexes of LRR-RPs, SOBIR1 and SERK proteins is lacking.

PRRs from various plant families have been shown to perceive small epitopes within larger microbial surface patterns that are sufficient to mediate plant immune activation^1–3^. Such immunogenic patterns are usually conserved among classes of microbes thus facilitating recognition of multiple microbial species through a single plant PRR. Furthermore, individual microbial patterns may have served as templates for the evolution of plant PRRs sensing structurally distinct immunogenic epitopes within these larger microbial patterns. A paradigm for this is bacterial flagellin perception in plants. FLS2, which recognizes an N-terminal 22-amino acid epitope (flg22) of bacterial flagellin^8^, is highly conserved in higher plants, including tomato^2,9^. However, tomato encodes an additional flagellin receptor, FLS3, which recognizes flagellin epitope flgII-28 that is distinct from flg22^10^. Likewise, rice harbors a yet unknown flagellin sensor that perceives a C-terminal epitope of *Acidovorax avenae* flagellin^11^, and *Vitis riparia* FLS2^XL^ senses *Agrobacterium tumefaciens* flg22^Atum^, a highly diverged flg22 variant that is not recognized by FLS2^12^. Analogous to flagellin sensing, *Brassicaceae-*specific ELONGATION FACTOR-THERMOUNSTABLE (*EF*-Tu) RECEPTOR (EFR) recognizes a conserved N-terminal N-acetylated epitope (elf18), whereas a central fragment of *EF*-Tu (EF50) is perceived by a yet unknown receptor in rice^13^.

Previous work has revealed that fungal endopolygalacturonases (PGs) are recognized by *A. thaliana* (At) RLP42/RBPG1 (RESPONSIVENESS TO BOTRYTIS POLYGALACTURONASES 1)^14^. In this study, we identify fungal PG fragment pg9(At), consisting of 9 amino acids, that is sufficient to activate immune responses and disease resistance in *A. thaliana* in a SOBIR1/SERK-dependent manner. We further show that closely related *A. arenosa* (Aa) does not respond to pg9(At), but to pg20(Aa), which is structurally distinct from pg9(At). Likewise, *Brassica rapa* (Bra) senses the fungal PG epitope pg36(Bra) but not pg9(At) or pg20(Aa). Our study sheds light on the complexity of PG pattern recognition receptor evolution in closely related *Brassicaceae* species.

## Results

### Identification of an A. thaliana defense-stimulating minimal structural motif within microbe-derived PGs

Most PRRs identified recognize immunogenic fragments within larger microbial surface signatures. Immunogenic activities of PG3 and PG6 from *Botrytis cinerea* are recalcitrant to reduction by dithiothreitol (DTT) and to digestion by endo proteases Glu-C and Lys-C (Supplementary Fig.1a-c), suggesting that a PG-derived fragment rather than tertiary fold features determine PG elicitor activity. Protease digestion patterns revealed that elicitor activity may be contained within a large, central PG fragment produced upon endo protease Glu-C digestion or within two shorter, central PG fragments produced by endo protease Lys-C treatment (Fig. 1a). A 103-amino acid region covering these fragments was selected, and synthetic peptides spanning this region (pep1-4) (Fig. 1a) were tested as elicitors of ethylene production in *A. thaliana* leaves (Supplementary Fig. 1d). Pep3 triggered strong ethylene production, whereas all other peptides tested lacked activity (Fig. 1a and Supplementary Fig. 1d).

**Figure 1.**
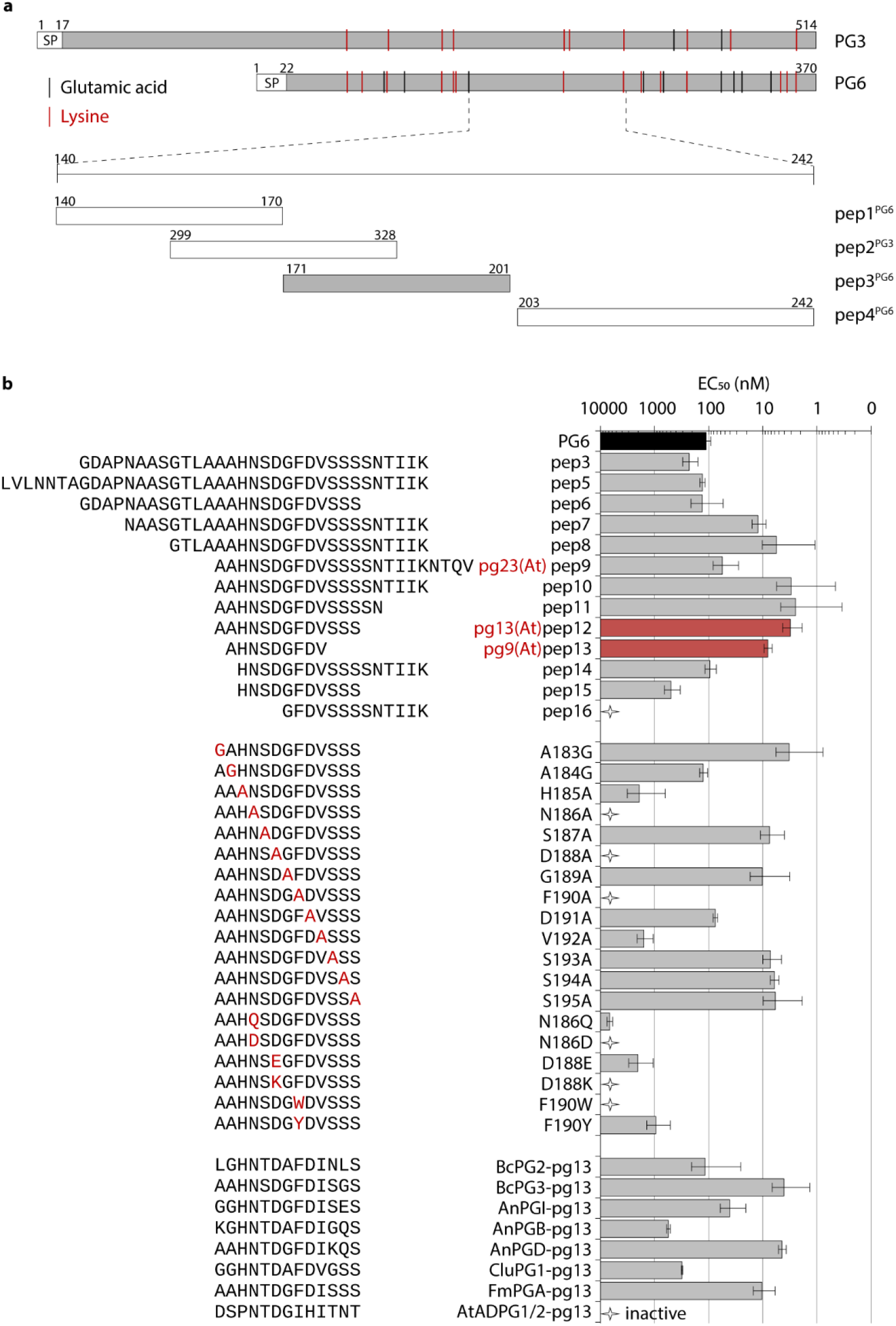
Identification of an *A. thaliana* defense-stimulating structural motif in *Botrytis cinerea* polygalacturonase 6. **a**, Schematic representation of PG gene sequence fragments. Glutamic acid and lysine residues within full-length PG3 and PG6 proteins are indicated as black and red lines, respectively. Numbers above the fragments indicate the start and end amino acid positions according to the corresponding PG sequence. Synthetic peptides pep1 to pep4 are derived from PG3 or PG6 are indicated. Protein/peptides are represented in grey or white depending on their ability or inability to induce ethylene production, respectively. Synthetic Pep2 derived from PG3 was tested because the corresponding peptide from PG6 was not obtained. **b**, Ethylene-inducing activity in *A. thaliana* Col-0 of peptides derived from fungal PGs. EC_50_ values were determined from dose-response curves with the synthetic peptides. In the right panel, bars represent means ± standard deviation on a logarithmic scale from at least two independent biological repeats, each with three replicates. The peptides that did not induce any ethylene production at up to 10 μM are defined as inactive peptide (stars). The peptides derived from various fungal PGs include BcPG2 and BcPG3 from *Botrytis cinerea*; AnPGI, AnPGB, and AnPGD from *Aspergillus niger*; CluGP1 from *Colletotrichum lupine*; and FmPGA from *Fusarium moniliforme*. AtADPG1/2-pg13 is derived from AtADPG1/2 of *A. thaliana*.

To delineate the minimal structural requirements for elicitor activity, a set of nested synthetic peptides spanning pep3 was assessed for elicitor activity (Fig. 1b). Among these, the 13-amino acid peptide pep12 was over 30-fold more active than full length PG6 (EC_50_ values of 3.1 and 111.7 nM, respectively) (Fig. 1b). By contrast, peptides with longer N- or C-terminal ends had similar activity to the full-length protein. The highly active pep12 was renamed pg13(At) (Fig. 1b).

To identify the amino acids essential for the activity of pg13(At), alanine-scanning mutagenesis (except for A183G and A184G) was conducted (Fig. 1b). Replacements of residues A184, H185, D191, and V192 led to 20- to 500-fold lower activities; whereas replacements of residues N186, D188, and F190 resulted in inactive peptides (Fig. 1b). Notably, D188 is a conserved PG residue required for catalytic activity^15^. Residues N186, D188, and F190 in pg13(At) were further mutagenized to chemically similar or dissimilar amino acids. These mutations also abolished or significantly reduced immunogenic activity (Fig. 1b). As residues A183 and S193-195 are dispensable for immunogenic activity (Fig. 1b), an additional synthetic 9-amino acid-peptide pg9(At) was generated and analyzed. Pg9(At) had a similar activity to pg13(At) (EC_50_ value of 8.1 nM, Fig. 1b), and thus the two peptides were used interchangeably in further experiments. Inactive analogs derived from elicitors may act as competitive antagonists of the corresponding elicitors to block or dampen the immune responses^16–18^. However, replacements of residues N186, D188, and F190 exhibited no antagonistic effect on ethylene accumulation upon pg9(At) elicitation (Supplementary Fig. 1e).

PG-encoding gene sequences are found in fungal, bacterial, oomycete and plant genomes (CAZy, http://www.cazy.org/Genomes.html). Synthetic pg13(At)/pg9(At) peptides derived from other fungi, bacteria, oomycete, and *A. thaliana* were thus analyzed for their immunogenic potential. *B. cinerea*-derived peptides as well as those from *Aspergillus niger*, *Colletotrichum lupine* and *Fusarium moniliforme* were active elicitors albeit with different EC_50_ values (Fig. 1b). Pg9(At) derived from PGs of *Phytophthora sp.* induced residual ethylene production with an EC_50_ value of 1.3 μM, which was ~160 times less active than pg9(At). In contrast to fungal pg13(At) peptides, pg13(At)/pg9(At)-like peptides derived from *A. thaliana* and *Xanthomonas sp.* exhibited no activity, demonstrating the specificity for fungal PG recognition (Fig. 1b and Supplementary Fig. 1f).

### Pg9(At)/pg13(At) responses are mediated by RLP42

*A. thaliana* accession Br-0 is unresponsive to PGs due to the absence of RLP42^14^. Pg9(At) failed to induce ethylene production in Br-0, but did so in a Br-0 line overexpressing RLP42 (Br-0+RLP42) (Fig. 2a). In addition to ethylene production, pg13(At)/pg9(At) triggered ROS burst, MAPK activation, and expression of the defense-related genes *FRK1* and *PAD3* in Col-0 and Br-0+RLP42, whereas accession Br-0 was unresponsive (Supplementary Fig. 2). To determine whether pg9(At) is sufficient to induce disease resistance, Br-0 and Br-0+RLP42 leaves were infiltrated with pg9(At) one day prior to inoculation with virulent *Pseudomonas syringae* pv. *tomato* (*Pst*) DC3000. Pg9(At) pre-treatment restricted bacterial growth on Br-0+RLP42 to a similar extent as nlp20 treatment, but failed to restrict bacterial infection in Br-0 (Fig. 2b). Unlike other known PAMPs (e.g. flg22, nlp20, chitin), PGs trigger hypersensitive response (HR)-like cell death on *A. thaliana*^14^. Many pep3-derived peptides, including the 23-amino acid pep9, also consistently induced cell death (Fig. 1b and Supplementary Fig. 3a). Pep 9 was renamed pg23(At) (Fig. 1b and Supplementary Fig. 3a). Unlike pg23(At) and longer peptides, leaves infiltrated a single time with pg9(At) and pg13(At) did not induce consistent cell death (Fig. 2c and Supplementary Fig. 3a). We hypothesized that this was due to instability or a short half-life of the shorter peptides. In support of this hypothesis, infiltration of Col-0 leaves with pg9(At) three times (at 0, 24, and 48 h) induced cell death similar to infiltration a single time with pg23(At) (Fig. 2c). A pg23(At) variant carrying mutations in residues required for RLP42 recognition (pg23m1) failed to induce cell death, whereas a variant carrying mutations outside the core pg9(At)-epitope retained its ability to induce both ethylene production and cell death (Supplementary Fig. 3b). Collectively, these results demonstrate that peptides containing the minimal pg9(At)-epitope activate plant immune responses in an RLP42-dependent manner.

**Figure 2.**
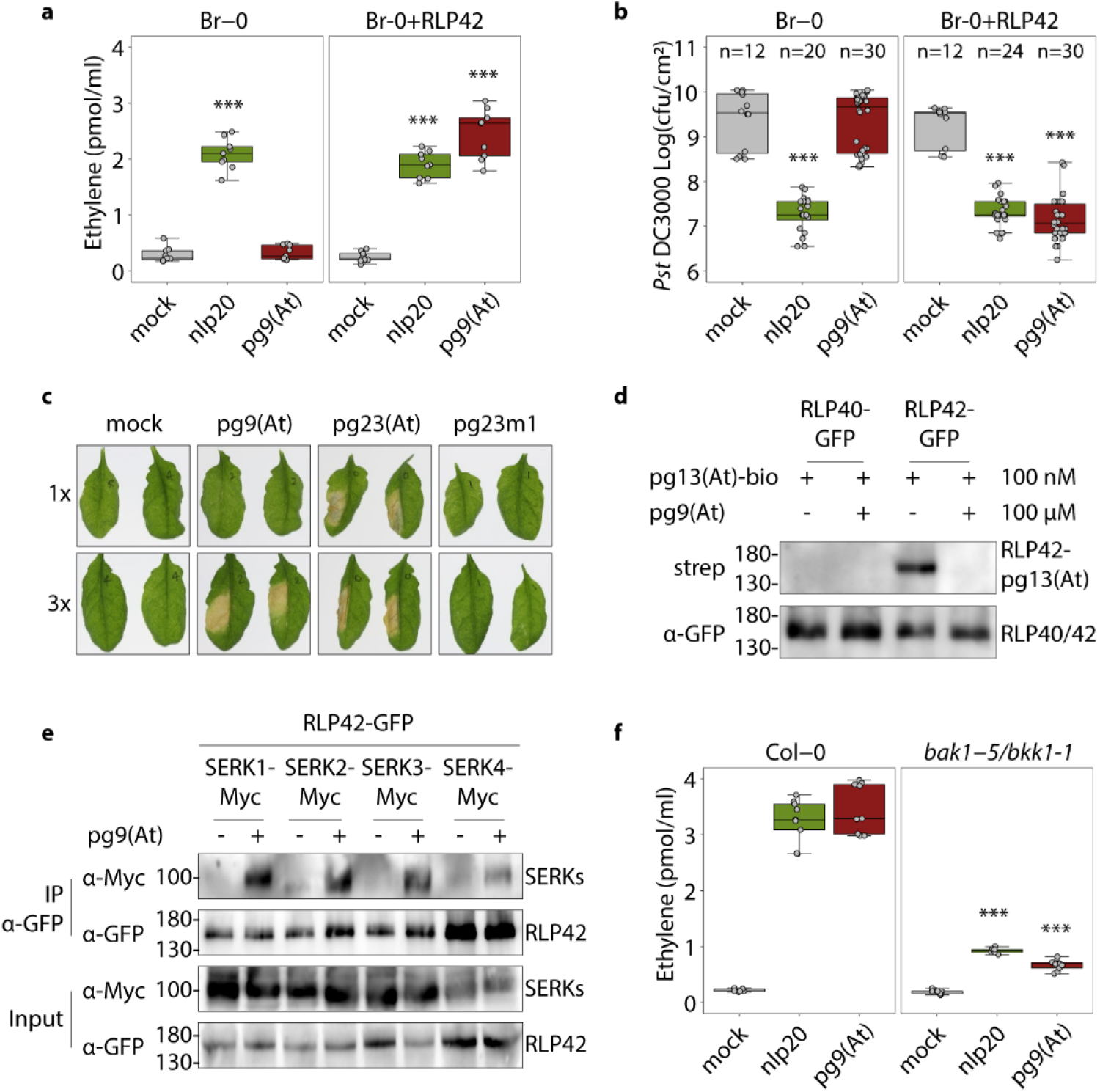
RLP42 binds to pg9(At) and forms a complex with SERK family proteins to activate plant immunity. **a**, Ethylene accumulation after 4 h treatment with 1 μM nlp20, pg9(At), or water (mock) in Br-0 and Br-0+RLP42 (n=9). **b**, Bacterial growth in Br-0 and Br-0+RLP42 plants treated with water (mock), or 1 μM nlp20 or pg9(At) 24 h prior to infiltration of *Pst* DC3000. Bacteria were quantified in extracts of leaves at 2 days post-inoculation. Results are representative of three independent experiments. **c**, Hypersensitive-like cell death in Col-0 leaves infiltrated with 10 μM of pg9(At), pg23(At), or pg23m1, or water (mock) once (1x) or every 24 h up to 3 times (3x) and visualized at 7 days post first infiltration. **d**, RLP42, but not RLP40, specifically binds pg13(At). Ligand binding assay using *Nicotiana benthamiana* transiently expressing RLP42-GFP as receptor or RLP40-GFP as negative control, biotinylated pg13(At) [pg13(At)-bio] peptides as ligand, and unlabelled pg9(At) as competitor. Immunoblot analysis using streptavidin-AP shows the presence of RLP42-pg13(At)-bio complex (top panel), and anti-GFP antibody shows the presence of RLP42-GFP in all samples (bottom panel). **e**, Proteins extracted from *N. benthamiana* leaves expressing RLP42-GFP in combination with Myc-tagged SERK family proteins and treated with water (−) or 1 μM pg9(At) (+) for 5 min before harvesting, were used for co-immunoprecipitation with GFP-trap beads, and immunoblotting with tag-specific antibodies. **f**, ethylene accumulation after 4 h treatment with 1 μM pg9(At), or water and 1 μM nlp20 as controls in A. thaliana Col-0 and *bak1-5/bkk1-1* mutant (n=9). For **a**, **b**, and **f**, data points are indicated as grey dots and plotted as box plots. Asterisks for **a** and **b** indicate statistical differences to mock treatments in the respective plant; for **f** asterisks indicate a significant difference between the mutant and Col-0 response for the given elicitor (*** *P*<0.001, Student’s *t*-test).

To determine whether pg13(At) binds RLP42 for immune activation, ligand binding assays were performed using immunogenically active (Supplementary Fig. 4) biotinylated pg13(At) [pg13(At)-bio]. *Nicotiana benthamiana* leaves expressing RLP42-GFP or RLP40-GFP (unresponsive to PG^14^) were treated with pg13(At)-bio and a chemical cross-linker. Subsequently, GFP-tagged proteins were immunoprecipitated using GFP affinity beads and detected by both GFP- and streptavidin-antibodies. Pg13(At)-bio was detected in RLP42- but not RLP40 pull downs (Fig. 2d). Addition of a large excess of free pg9(At) competitively abolished pg13(At)-bio binding, demonstrating that RLP42 specifically binds pg13(At) (Fig. 2d).

We next asked whether elicitor binding induces recruitment of SERK family co-receptors. We found that RLP42-GFP forms a complex with four SERK family proteins (SERK1, SERK2, SERK3/BAK1, and SERK4/BKK1) in a pg9(At)-dependent manner (Fig. 2e). Consistent with this finding, a *bak1-5/bkk1-1* genotype was strongly impaired in response to pg9(At) and pg23(At) (Fig. 2f and Supplementary Fig. 5). Together with previous data showing that the constitutively associated SOBIR1 is required for RLP42 signaling^14^, these data demonstrate that elicitor binding by RLP42 leads to the formation of a tri-partite active signaling complex, analagous to what has been reported for RLP23^19^.

### Structure/function analysis of RLP42-ligand interaction

To further investigate the molecular basis of RLP42 function, we performed a structure-function analysis of RLP42 based on sequence analysis of RLP42 and its closely related paralogs. RLP39 to RLP42 are four paralogs with over 80% amino acid identity to each other, but only RLP42 is responsive to PGs^14^. Sequences of RLP42 and RLP40 were used to design domain swap constructs. Both proteins have an extracellular domain with 25 LRRs interrupted by a 49-amino acid island domain (ID) (Fig. 3a). Six chimeric constructs (S1-S6) were generated consisting of RLP42 with a region replaced with the corresponding part from RLP40 (Fig. 3a). LRRs 13-15 were excluded for swapping as the amino acid sequences are identical between RLP42 and RLP40. The chimeric proteins were transiently expressed in *N. benthamiana* at similar levels (Supplementary Fig. 6a) and assessed for their ability to perceive pg9(At). Upon pg9(At) elicitation, the levels of ROS burst and ethylene production were similar among leaves expressing RLP42, S4, and S6; whereas the levels were significantly reduced in leaves expressing S1, S2, S3, and S5 (Fig. 3b and Supplementary Fig. 6b). Additionally, an ID deletion mutant (ΔID) exhibited no ethylene production upon pg9(At) treatment (Supplementary Fig. 6c). Collectively, these results demonstrate that the RLP42 N-terminus-LRR12 and LRR21-LRR24, including the ID, contain structural elements required for pg9(At) recognition.

**Figure 3.**
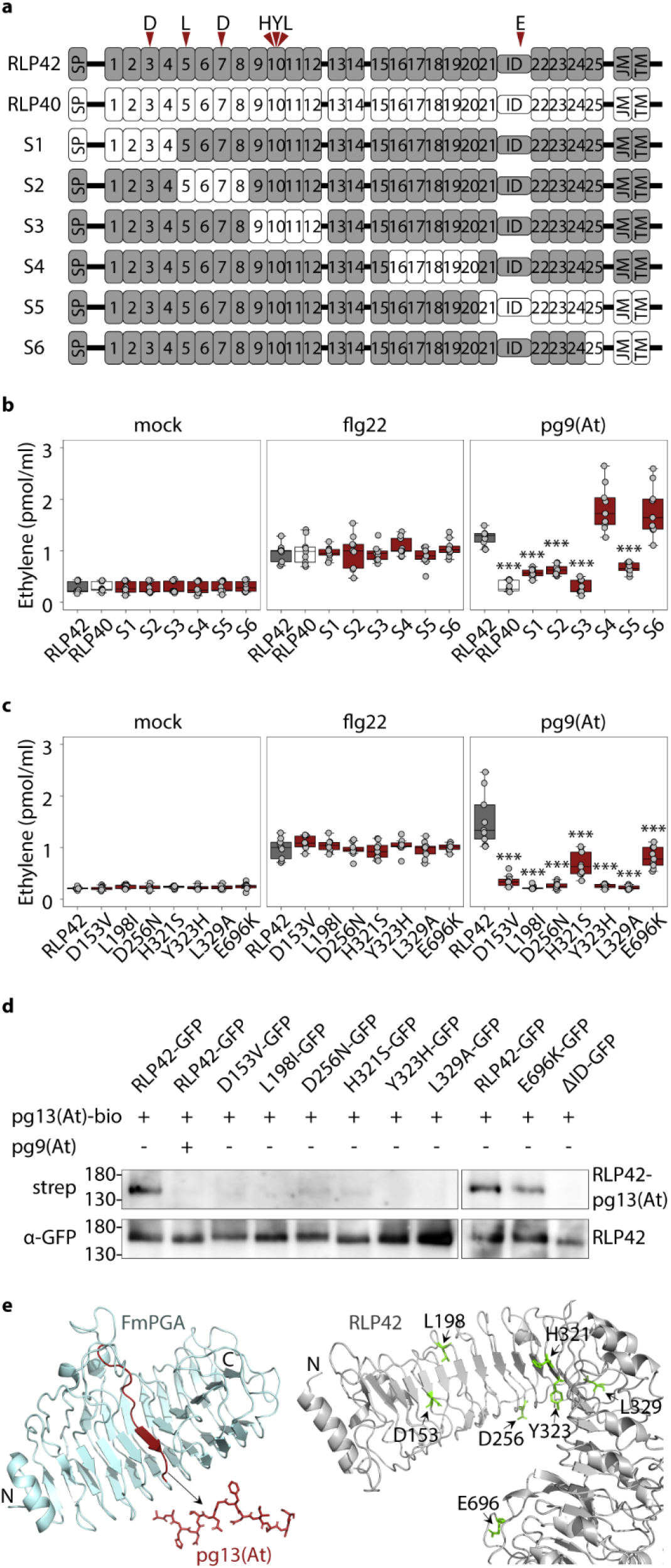
Structure-function analysis of RLP42. **a**, Schematic representation of the RLP42 and RLP40 chimeric proteins (S1-S6) used for structure-function analysis. SP (signal peptide), ID (island domain), JM (juxtamembrane domain), TM (transmembrane domain), and numbers indicate leucine-rich repeats. **b**,**c**, Ethylene production after 4 h treatment with 1 μM pg9(At) or flg22 or water (mock) in *N. benthamiana* leaves transiently transformed with the chimeric constructs (**b**) or the point mutation constructs (**c**). Data points are indicated as grey dots from three independent experiments (n=9) and plotted as box plots. Asterisks indicate significant differences to RLP42-expressing plant for the given elicitor (*** *P*<0.001, Student’s *t*-test). **d**, Binding of biotinylated pg13(At) [pg13(At)-bio] to RLP42 receptor mutant proteins. RLP42/mutant-GFP transiently expressed in *N. benthamiana* as receptor, pg13(At)-bio peptides as ligand, and unlabelled pg9(At) as competitor. Immunoblot analysis using streptavidin-AP shows the presence/absence of RLP42-pg13(At)-bio complex (top panel), and anti-GFP antibody shows the presence of RLP42-GFP in all samples (bottom panel). The binding assay was performed in triplicate with similar results. **e**, Structural analysis of pg13(At) and RLP42 by PyMOL. Location of pg13(At) within *Fusarium moniliforme* PGA (PDB ID: 1HG8) is highlighted in red (left panel). A homology model of RLP42 was generated by SWISS-Model based on the BRI1 crystal structure (PDB ID: 3RGX). The important amino acids for pg13(At) perception are highlighted in green.

To further pinpoint amino acids required for pg9(At) recognition, differences of single amino acids or short stretches (2-10 amino acids) in extracellular domains of RLP39-42 were manually inspected. In total, 25 mutations were made in RLP42, and the proteins were transiently expressed in *N. benthamiana* (Supplementary Fig. 6). Variants with mutations in LRR4 (D153V), LRR5 (L198I), LRR7 (D256N), LRR10 (H321S, Y323H, L329A) and in the ID (E696K) were impaired in ethylene production and ROS burst in response to pg9(At) (Fig. 3c and Supplementary Fig. 6b, d). All mutants, except H321S and E696K, were also impaired or significantly reduced in pg13(At)-bio binding (Fig. 3d) and elicitor-induced BAK1 recruitment (Supplementary Fig. 6e). Likewise, the ΔID mutant exhibited no pg13(At)-bio binding and was impaired in pg9(At)-induced BAK1-recruitment. Collectively, these data highlight LRR4, −5, −7, and −10 and the ID as important structural features for RLP42 function (Fig. 3e).

### Pg13(At) perception is phylogenetically restricted

To assess the distribution of the pg13(At) recognition system among plants, 52 *A. thaliana* accessions were tested for pg13(At)-induced ethylene production. Unlike other LRR-RP patterns (e.g. eMax, nlp20, SCFE1, and IF1) that are perceived by most of the accessions studied^19–23^, pg13(At) was perceived by only 16 of the 52 accessions (Fig. 4a and Supplementary Fig. 7a). The responsiveness to pg13(At) was further tested in 16 additional plant species. Surprisingly, only *A. thaliana* was responsive to pg13(At) (Supplementary Fig. 7b), whereas most of the plants tested were responsive to flg22 and several *Brassicaceae* species were responsive to nlp20 (Supplementary Fig. 7b), consistent with previous findings^24,25^. These results indicate that the pg13(At) recognition system is restricted in *A. thaliana* with notable accession specificity.

**Figure 4.**
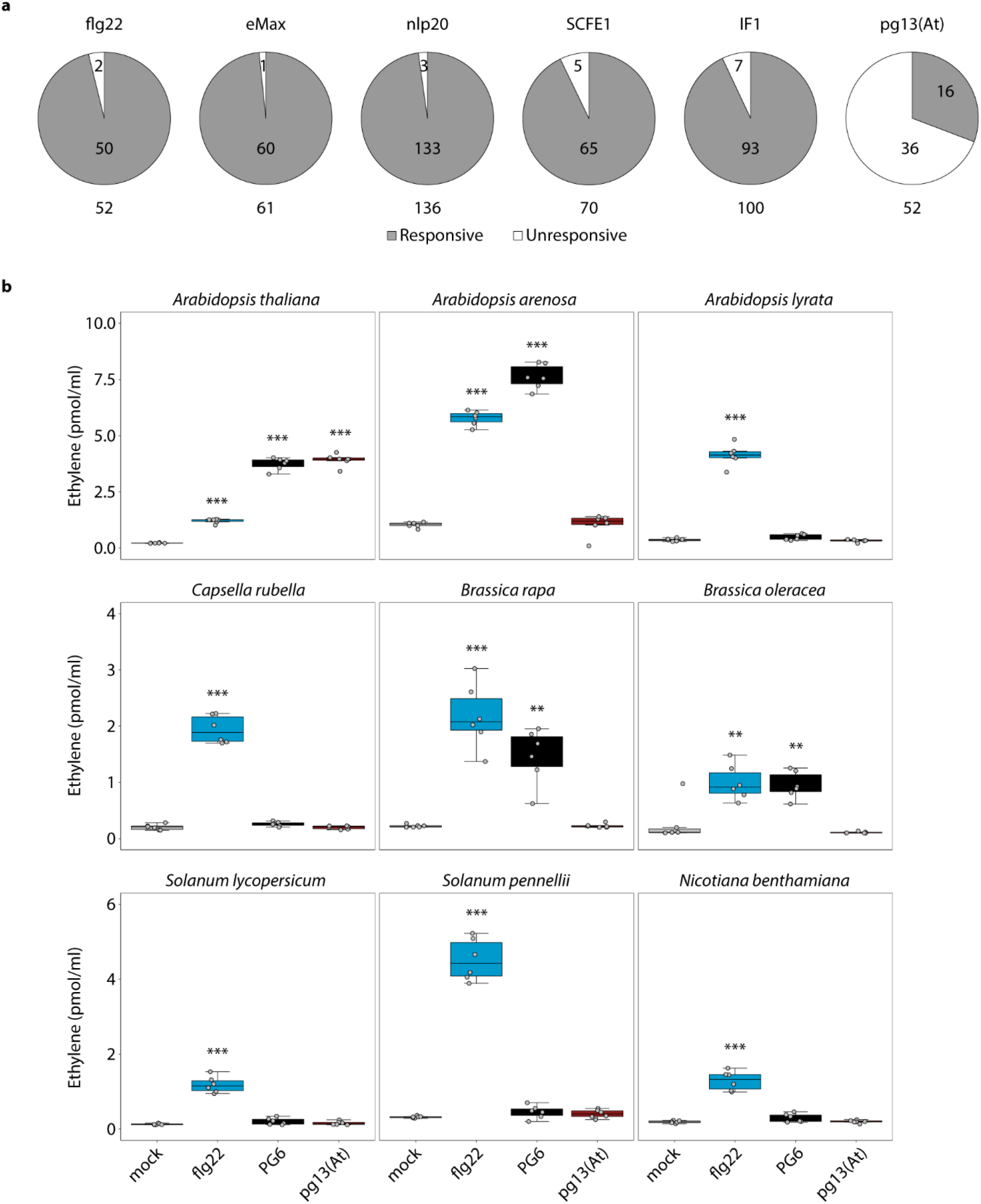
Immunogenic activity of PG6 and pg13(At) in *A. thaliana* accessions and in various plant species. **a**, 52 *A. thaliana* accessions were tested for the ethylene accumulation upon 1 μM pg13(At) or flg22, or water (mock) treatment. An increase of >2-fold relative to mock treatment was scored as responsive. Data for eMax, nlp20, SCFE1, and IF1 were obtained from the literature^19–23^. Numbers in pie charts indicate the number of accessions in each category. Numbers below pie charts indicate the total accessions studied. **b**, Ethylene accumulation after 4 h treatment with water (mock) or 50 nM PG6, 1 μM flg22 or pg13(At). Data points, indicated as grey dots, are from two independent experiments (n=6) and plotted as box plots. Asterisks indicate statistical differences to mock treatments in respective plant (*** *P*<0.001, ** *P*<0.01, Student’s *t*-test).

### Identification of additional PG motifs with immunogenic activity in Brassicaceae

Although insensitive to pg13(At), several *Brassicaceae* species were responsive to full-length PG6 protein (Fig. 4b) suggesting the presence of additional immunogenic motif(s) within PG6. To test this idea, the immunogenic activities of peptides pep1-4 (Fig. 5a) were reassessed in *A. arenosa* and *B. rapa.* Pep1-3 were inactive (Supplementary Fig. 8), but pep4, which does not contain pg13(At), induced ethylene production in both plants (Fig. 5b, Supplementary Fig. 8). To identify the minimal immunogenic epitope for activation of immunity in these species, a set of nested synthetic peptides from PG6 spanning pep4, named sequentially pep17-24, were analyzed for their ability to trigger ethylene production (Fig. 5b). In *A. arenosa*, a 20-amino acid peptide was found to be as active as pep4 and was renamed pg20(Aa) (Fig. 5b). However, pg20(Aa) was only weakly active in *B. rapa* (EC_50_ value of 2,259 nM). The longer peptide pep22 [hereafter pg36(Bra)] was the most active of the peptides tested in *B. rapa* with an EC_50_ value of 181 nM (Fig. 5b).

**Figure 5.**
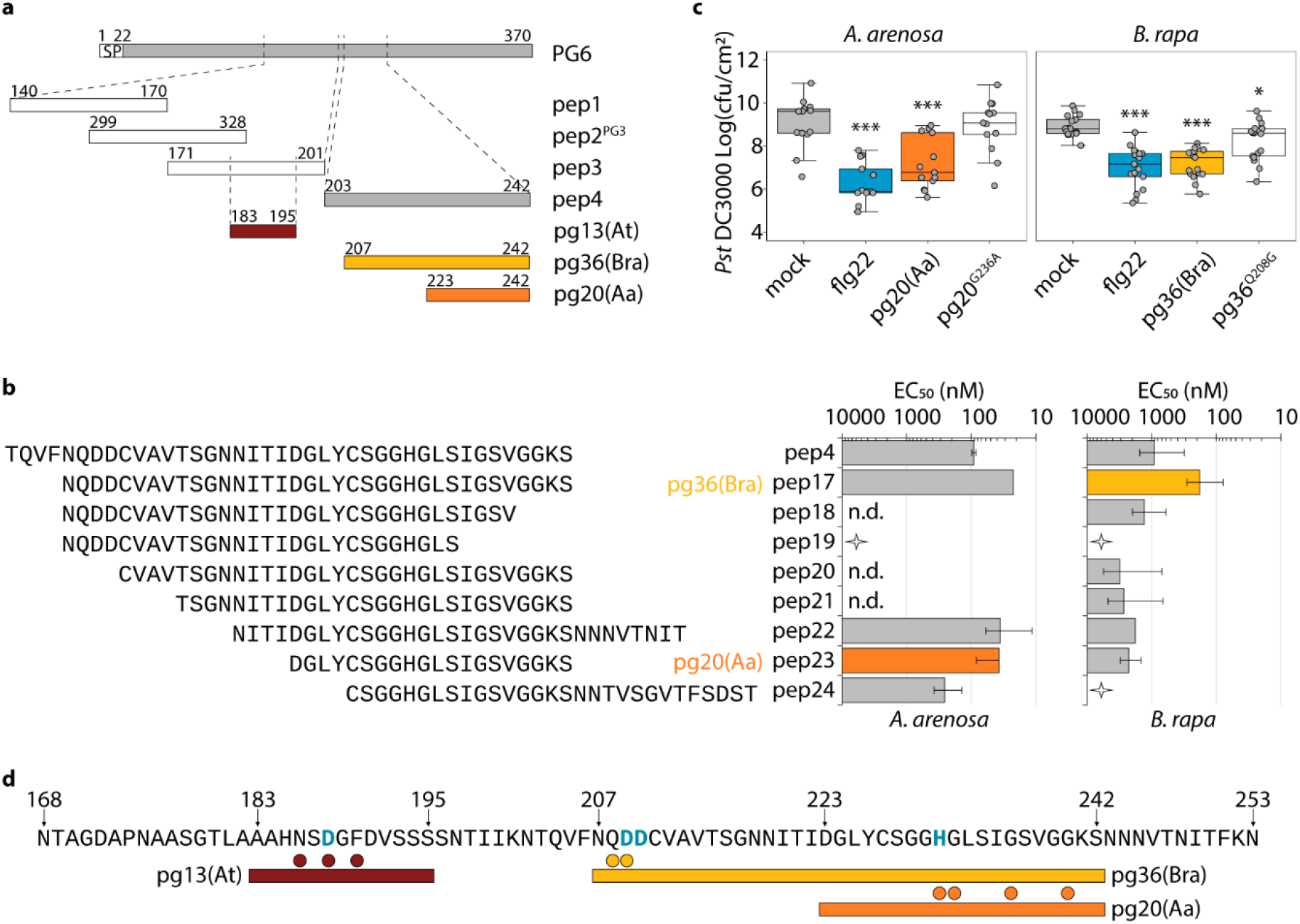
Identification of PG epitopes triggering immunity in *A. arenosa* and *Brassica rapa*. **a**, Schematic representation of protein PG6 and the peptides pep1-4 used for pg20(Aa) and pg36(Bra) identification. Numbers above the peptides indicate the start and end amino acid positions according to PG6 sequence, except for the positions of pep2, which indicate positions of the PG3 sequence. **b**, Ethylene-inducing activity of peptides derived from PG6 in *A. arenosa* and *B. rapa*. EC_50_ values were determined from dose-response curves with the synthetic peptides. Bars represent means ± standard deviation on a logarithmic scale from at least three independent biological repeats, each with three replicates. The peptides that did not induce any ethylene production at up to 10 μM are indicated with stars. n.d., not determined. **c**, Bacterial growth in *A. arenosa* and *B. rapa* plants treated with water (mock), or 1 μM the given peptide 24 h prior to infiltration of *Pst* DC3000. Bacteria were quantified in extracts of leaves at 4 (*A. arenosa*) or 2 (*B. rapa*) days post-inoculation. Data points are indicated as grey dots from three independent experiments (for *A. arenosa*, n=13; for *B. rapa*, n=17) and plotted as box plots. Asterisks indicate significant differences to mock treatments in the respective plant (*** *P*<0.001, * *P*<0.05, Student’s *t*-test). **d**, Schematic representation of the sequence and position of pg13(At) (red bar), pg20(Aa) (orange bar), and pg36(Bra) (yellow bar) within PG6. The colored dots above the bars indicate the important residues required for recognition in the corresponding peptides. The residues required for PG6 enzymatic activity are indicated in blue. Numbers above the residues indicate the positions according to the PG6 sequence.

Alanine-scanning mutagenesis was performed to identify the amino acids essential for the activity of pg20(Aa) in *A. arenosa* (Supplementary Fig. 9). Replacements of residues H231, G232, G236, and G240 led to only residual activity (EC_50_ > 1 μM) (Supplementary Fig. 9), indicating the importance of these residues in pg20(Aa) perception. Notably, H231 is also required for fungal PG enzymatic activity^15^. Similarly, analysis of point-mutated pg36(At) revealed that Q208 and D209 are important for perception in *B. rapa* (Supplementary Fig. 10), with residue D209 being conserved and required for fungal PG enzymatic activity^15^. Pg36(Bra)-like peptides derived from *B. rapa* (Bra-pg36) itself exhibited no activity (Supplementary Fig. 10). Pg20(Aa) and pg36(Bra) pre-treatment enhanced resistance to the bacterial pathogen *Pst* DC3000 in *A. arenosa* and *B. rapa*, respectively; whereas unrecognized variants pg20^G236A^ and pg36^Q208G^ failed to confer pathogen resistance (Fig. 5c). Altogether, these results demonstrate that *A. arenosa* and *B. rapa* perceive overlapping PG epitopes that are distinct from pg13(At)/pg9(At) recognized by RLP42 (Fig. 5d). Requirement of PG residues Q208 and D209 for pg36(Bra) (Fig. 5d and Supplementary Fig. 10), but not for pg20(Aa) elicitor activity further suggests that PG sensor systems in *A. arenosa* and *B. rapa* exhibit different ligand binding specificities.

## Discussion

In a previous study, constitutively formed *A. thaliana* RLP42 SOBIR1 complexes have been implicated in the activation of fungal PG-inducible plant defenses, including hypersensitive cell death^14^. In this study, we identified and characterized a minimal epitope pg9(At) from fungal PGs that is sufficient to bind to the receptor RLP42 and to trigger immunity in *A. thaliana*. Pg9(At) peptides from various fungal PGs, but not those from bacterial, oomycete or plant PGs are efficiently sensed by RLP42, suggesting that this receptor may facilitate recognition of a large number of fungal species. Pg9(At) comprises part of the core beta barrel structure of PG (Fig. 3e) and makes up part of the substrate binding cleft^15,26–28^. Such sequences are often evolutionarily stable with low levels of polymorphism, which may explain why such motifs have become preferred templates for PRR evolution. We further show that ligand binding recruits functionally redundant SERK protein family members into a ternary RLP42-SOBIR1-SERK complex. In sum, RLP42 shares with other LRR-RP-type PRRs similar characteristics of ligand binding and receptor complex assembly.

X-ray crystallography analysis of the FLS2-BAK1 complex with flg22 revealed that LRRs 3-16 of FLS2 contribute to flg22 binding. Flg22 interacts with both FLS2 and BAK1, acting as a molecular glue to promote the association of receptor and co-receptor^7^. Despite intense attempts by several labs, structural elucidation of any plant LRR-RP complex has thus far been unsuccessful. Thus, how this class of receptors interacts with their ligands and co-receptors remains unclear. Limited insight into that has recently resulted from structure-function analysis with LRR-RP RLP23, which revealed the importance of N-terminal LRRs 1-3 for ligand binding^29^.

The RLP42-pg9(At) system presents a useful new model for characterization of LRR-RP function because RLP42 has 3 closely related, but inactive paralogs, thus allowing for more detailed insights into structural requirements of ligand-receptor interaction and PRR complex assembly. Our structure-function analyses revealed seven residues of RLP42 that are crucial for pg9(At) perception, of which six are located in LRR3-10 (Fig. 3e) and one in the island domain (ID). We also found that a ΔID mutant is unable to bind the ligand and recruit BAK1 to the signal complex. Pg9(At) is a rather small ligand; we hypothesize that pg9 binds to the N-terminal LRRs (LRR3-10) of RLP42 and stabilizes the island domain of RLP42 for BAK1 recruitment. Unlike flg22 and brassinosteroids, which contact both the receptor (FLS2 and BRI, respectively) and the coreceptor BAK1^7,30,31^, pg9(At) might function similar to plant growth factor phytosulfokine (PSK), a tyrosine-sulfated pentapeptide. PSK binds to the island domain of the PSK-receptor (PSKR) thereby stabilizing this domain and, subsequently, facilitating SERK co-receptor recruitment^32^.

RLP42 is phylogenetically restricted. Pg13(At) responsiveness was only detected in *A. thaliana,* and less than a third of *A. thaliana* accessions tested were sensitive to pg13(At). This extent of accession-specific sequence polymorphisms or scattered distribution of functional PRR alleles is rather reminiscent of that reported for intracellular nucleotide-binding LRR proteins (NLRs) mediating ETI, and distinguishes RLP42 from most other PRRs (Fig. 4a). In a recent study, we have shown that *A. thaliana* LRR-RP genes share with NLRs not only a genomic organization into gene clusters but also apparently similar evolutionary dynamics maintaining large sequence diversity, while LRR-RK-encoding genes are much more uniform^6^. Altogether, these findings imply that evolution of LRR-RP-type PRRs is much more rapid than reflected in prevailing models of plant immunity, that suggest substantially different rates underlying the evolution of PRRs and NLRs^4^.

This notion is further supported by the fact that *A. thaliana*, *A. arenosa*, and *B. rapa* recognize different immunogenic epitopes within PGs. Whereas pg9(At)/pg13(At), is recognized by RLP42 in *A. thaliana*, PG perception system(s) in *A. arenosa* and *B. rapa* are yet to be identified. However, as pg9(At) and pg20(Aa) are physically separate parts within PG (Fig. 5d), it is reasonable to assume that *Brassicaceae* have evolved at least two different perception systems for fungal PGs.

The pg20(Aa) epitope is contained entirely within pg36(Bra) (Fig. 5d). Residues Q208 and D209 of pg36(Bra) are required for elicitor activity in *B. rapa* (Supplementary Fig. 10), but are not part of pg20(Aa), which triggers immunity in *A. arenosa* (Fig. 5b). Likewise, residue G240, which is part of both fragments, is required for elicitor activity of pg20(Aa), but not for that of pg36(Bra) (Fig. 5d, Supplementary Fig. 9, 10). This may suggest that the recognition of these two epitopes is brought about by independently evolved receptors in the two plants. However, this scenario may also be explained by different alleles of the same receptor exhibiting different ligand specificities. Two alleles of FLS2, from *A. thaliana* and tomato, both bind flg22, but the tomato FLS2 is also capable of binding flg15, a shorter fragment of the same epitope^33^. Identification of the cognate receptor(s) in *B. rapa* and *A. arenosa* is required to address whether closely related *Brassicaceae* species harbor three distinct perception systems for PG-derived fragments.

Recognition of different fragments within larger immunogenic patterns through distinct PRRs has been reported before. So far, however, such examples relate to species only from remotely related plant families. Our findings now imply that evolution of distinct sensor systems for the same microbial pattern can be observed as well in closely related species within the same plant family. Our insights further suggest that some plant PRRs may share with NLR sensors similar evolutionary dynamics.

## Materials and Methods

### Plant materials and growth conditions

All plants were grown in soil and used at an age of 6-8 weeks. *A. thaliana* and *A. arenosa* plants were grown in a growth chamber at 22°C under short-day conditions of 8 h of light/16 h of dark. *Nicotiana benthamiana* and *Brassica* plants^34^ were grown in a greenhouse at 23°C under long-day conditions of 16 h of light/8 h of dark.

### Peptides

Synthetic peptides (GenScript) were prepared as 10 mM stock solutions in 100% dimethyl sulfoxide (DMSO), and diluted in water to the desired concentration prior to use.

### Plant immune responses

Reactive oxygen species (ROS) burst, ethylene production, MAPK activity assay, and gene expression analysis were performed as described^35^. For ROS burst, leaf pieces were floated in 96-well plates (1 piece well^−1^) containing 100 μl of solution with 20 μM L-012 (Waco) and 20 ng ml^−1^ horseradish peroxidase (Applichem); Luminescence after treatment with peptides or water (as control) was measured with a luminometer (Mithras LB 940, Berthold) in 2 min intervals. Total relative light unit (RLU) production was determined by calculating the area under the scatter curve for the time points indicated. The RLU values at time point 0 min were set as 0 to accordingly remove the background responses. For ethylene production, three leaf pieces were incubated in a sealed 6.5 ml glass tube with 0.4 ml 20 mM MES buffer, pH 5.7 and the indicated elicitors. Ethylene production was measured by gas chromatographic analysis (GC-14A; Shimadzu) of 1 ml air from the closed tube after 4 h incubation. MAPK activity assay was performed by immunoblotting with anti-phospho p44/42 MAP kinase antibody (Cell Signaling Technology). For gene expression assays, infiltrated leaves were harvested after 6 h for total RNA isolation with NucleoSpin^®^ RNA Plus Kit (Macherey-Nagel). First strand cDNA was synthesized with RevertAid™ M-MuLV reverse transcriptase (Thermo Scientific) and qRT-PCR was performed with the iQ5 Multi-color real-time PCR detection system (Bio-Rad) using the SYBR Green Fluorescein Mix (Thermo Scientific) and gene-specific primers listed in Table S1. For cell death assays, leaves of *A. thaliana* were infiltrated with 10 μM pg23 or pg23m1, and leaves of *B. rapa* were infiltrated with 1 μM pg36 or pg36^Q208G^, and the absence or presence of chlorosis was observed after 7 days. For Fig. 2c, leaves were infiltrated either once or three times at 0, 24, and 48 h.

### Infection assay

Six- to 8-week-old plants were primed 24 h prior to infection by leaf infiltration of 1 μM peptide, or water (mock treatment). *Pseudomonas syringae* pv. *tomato* DC3000 was grown in King’s B medium at 28°C for ~16 h, and resuspended in 10 mM MgCl_2_ at a concentration of 5×10^5^, 1×10^6^, and 1×10^4^ colony-forming units (cfu) ml^−1^ for *A. thaliana*, *A. arenosa*, and *B. rapa* leaf infiltration, respectively. Bacterial populations were determined two or four days after infiltration.

### Construction of RLP42 chimeras and mutations

Domain swap constructs were generated using Gibson assembly Master Mix (New England BioLabs). The coding sequences of RLP42 and RLP40 were cloned into pDONR207 (Invitrogen) and/or pLOCG vector^36^. Using the resulting plasmids as templates, the corresponding fragments were amplified with overlap regions and assembled with *Spe*I-digested pLOCG to generate the chimeric constructs with C-terminal GFP fusion. Mutation constructs were generated either using Gibson assembly Master Mix as described above or the GeneArt Site-Directed Mutagenesis System (Thermo Scientific) and AccuPrime Pfx DNA Polymerase (Thermo Scientific) using pDONR207::RLP42 as template. Mutated RLP42 sequences were recombined into pB7FWB2.0 (Plant Systems Biology, VIB, University of Ghent) for C-terminal GFP fusion. Primers are listed in Table S1.

### Transient expression in N. benthamiana

*A. tumefaciens* (strain GV3101) harbouring the corresponding constructs were grown in LB medium with appropriate antibiotics at 28°C overnight, harvested and resuspended in 10 mM MgCl_2_, 10 mM MES pH 5.7, 200 μM acetosyringone to the desired OD600. The cultures carrying appropriate constructs were mixed to final OD600 = 0.1 per construct, incubated at room temperature for 2 h, and infiltrated into 6-week-old *N. benthamiana* plants. Leaves were cut into pieces 24 h after infiltration of agrobacteria, floated overnight on water, and used for ROS burst and ethylene production assays. For immunoprecipitation assays, leaves were harvested 40-48 h after infiltration of agrobacteria.

### In vivo cross-linking and immunoprecipitation assays

For *in vivo* cross-linking, leaves of *N. benthamiana* expressing RLP42-GFP were infiltrated with 100 nM pg13(At)-bio with or without 100 μM unlabelled pg9(At) as competitor. For cross-linking of peptides to the receptor, 2 mM ethylene glycol bis (succinimidyl succinate) [(EGS), initially dissolved in a small volume of DMSO and further diluted in 25 mM HEPES buffer pH 7.5] was co-infiltrated into the leaves. After 20 min of infiltration, the leaves were harvested and frozen in liquid nitrogen. For immunoprecipitations, membrane proteins of agro-infiltrated *N. benthamiana* leaves were extracted at 1 mg ml^−1^ in extraction buffer as described^14^ and immuno-adsorbed by means of their GFP-tags on GFP-Trap agarose beads (ChromoTek). Immunoblots were developed either directly with Streptavidin-alkaline phosphatase conjugate (Roche), or with anti-GFP antibodies (Torrey Pines Biolabs) or anti-myc antibodies (Sigma-Aldric), followed by staining with secondary antibodies coupled to alkaline phosphatase and CDP-Star (Roche) as substrates.

## Acknowledgments

This work was supported by Deutsche Forschungsgemeinschaft (DFG) grants Nu70/9-2, Nu 70/15-1, Nu70/16-1 and Nu70/17-1 to T.N. We thank Rebecca Schwab (Max Planck Institute for Developmental Biology, Tübingen) for providing *A. arenosa* seeds.

## Supporting information

**Supplementary Figure 1.**
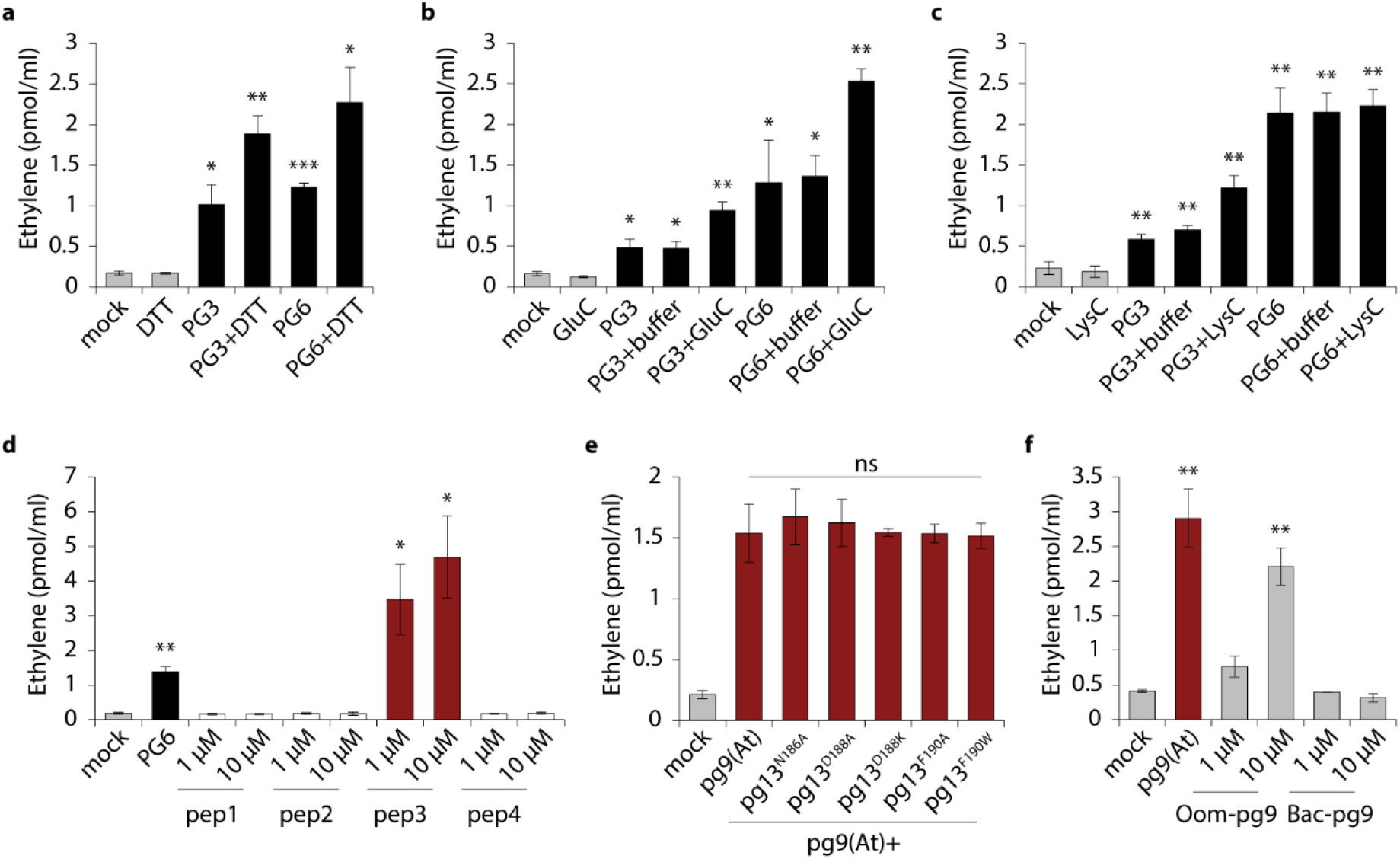
Ethylene-inducing activity of *Botrytis cinerea* PGs and peptides derived from PGs. Ethylene accumulation in Arabidopsis Col-0 was measured 4 hours after treatment with the following elicitors. **a,** 60 nM DTT treated/untreated PG3/PG6, or 5 mM DTT or water (mock) as negative control. **b,c,** 100 nM GluC (**b**) or LysC (**c**) treated/untreated PG3/PG6, or GluC/LysC or water (mock) as negative control. **d,** 50 nM PG6, or 1 or 10 μM synthetic peptides pep1-4, or water (mock) as negative control. **e**, 10 nM pg9(At) alone or together with 100 μM mutagenized pg13^N186A^, pg13^D188A^, pg13^D188K^, pg13^F190A^, or pg13^F190W^, or water (mock) as negative control. **f**, 1 μM pg9(At), Oom-pg9 (AKNTDGFDL, derived from PGs of *Phytophthora sp.*), or Bac-pg9 (AKNTDGFDP, derived from PGs of *Xanthomonas sp.*), or water (mock) as negative control. Bars represent means ± standard deviation of three replicates. Asterisks indicate significant differences to mock treatments (*** *P*<0.001, ** *P*<0.01, * *P*<0.05, Student’s *t*-test). ns, no significant difference to pg9(At) treatment.

**Supplementary Figure 2.**
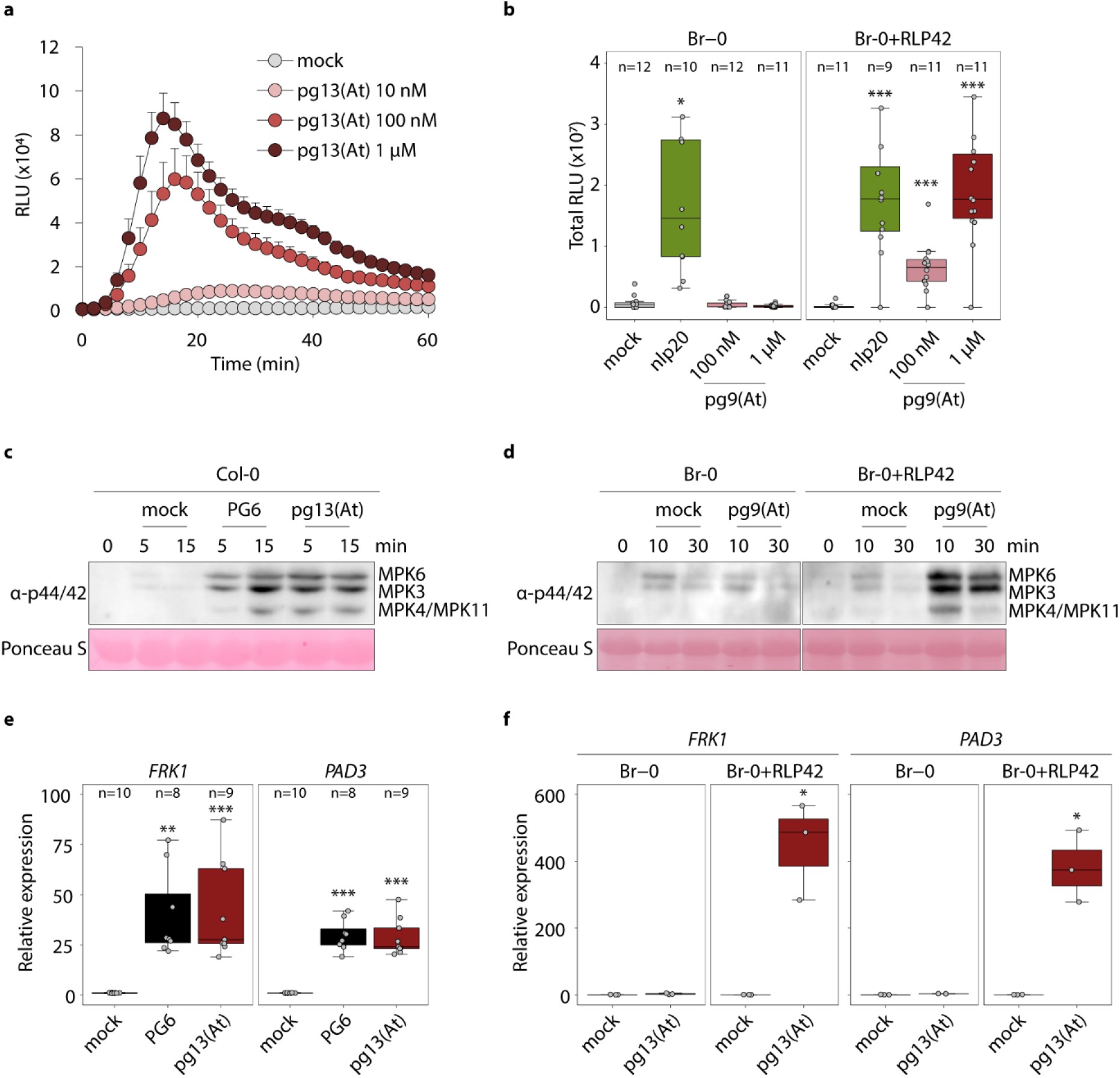
pg13(At)/pg9(At) activates plant immunity in RLP42-expressing Arabidopsis. **a,b,** Reactive oxygen species (ROS) production in Arabidopsis Col-0 (**a**) and Br-0 overexpressing RLP42 (Br-0+RLP42) or not expressing RLP42 (Br-0) (**b**) after treatment with water (mock) or the given elicitor. For **a**, bars represent means ± standard deviation (n=6) of relative light units (RLU). For **b**, total ROS production over 50 min were measured. **c,d,** Arabidopsis Col-0 (**c**) and Br-0 overexpressing RLP42 (Br-0+RLP42) or not expressing RLP42 (Br-0) (**d**) were infiltrated with water (mock) or 1 μM given elicitor and harvested at the indicated time points. Activated MAPKs were detected by immunoblot using anti-p44/42 MAP kinase antibody. Ponceau S stained blot is shown as a loading control. Assays were performed in triplicate with similar results. **e,f,** Transcript levels of *FRK1* and *PAD3* by quantitative real-time PCR (qRT-PCR). Arabidopsis Col-0 (**e**) and Br-0 overexpressing RLP42 (Br-0+RLP42) or not expressing RLP42 (Br-0) (**f**) were infiltrated with water (mock) or 1 μM given elicitor and sampled 6 h after treatment. Relative expression of the indicated genes was normalized to the levels of *EF-1α* transcript and calibrated to the levels of the mock treatment. For **b,e,f,** Data points are indicated as grey dots and plotted as box plots (**f**, n=3). Asterisks indicate significant differences to mock treatments in the respective plant (*** *P*<0.001, ** *P*<0.01, * *P*<0.05, Student’s *t*-test).

**Supplementary Figure 3.**
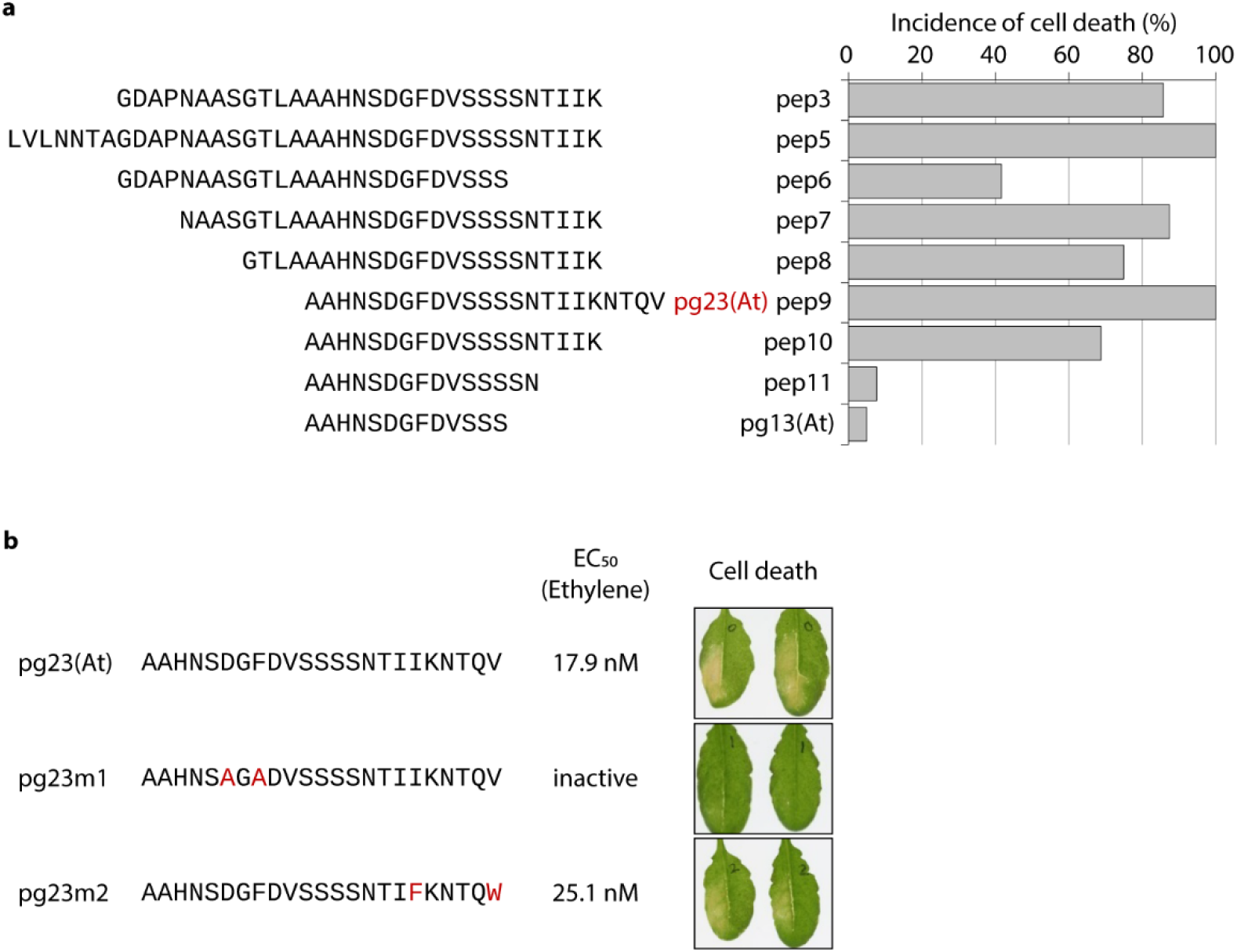
Cell death-inducing activity of peptides derived from PG6. **a,** Arabidopsis Col-0 leaves were infiltrated with 10 μM of the indicated peptides (left panel). Cell death symptoms were scored at 7 days post infiltration (dpi), and the incidence of cell death was calculated (right panel). **b,** The cell-death inducing activity of pg23(At) is associated with its immunogenic activity. The sequences of mutagenized peptides (pg23m1 and pg23m2) are indicated at the left panel. EC_50_ values were determined from dose-response curves with the synthetic peptides (middle panel). Leaves infiltrated with the indicated peptides were photographed at 7 dpi (right panel).

**Supplementary Figure 4.**
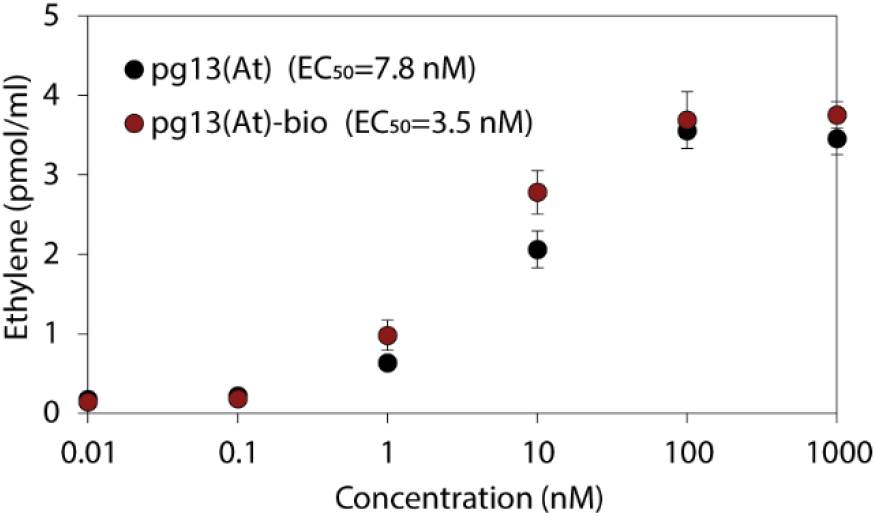
Biotinylated pg13(At) peptide is biologically active. Ethylene accumulation in Arabidopsis Col-0 was measured 4 hours after treatment with pg13(At), or biotinylated pg13(At) [pg13(At)-bio]. EC_50_ values were determined from dose-response curves. Bars represent means ± standard deviation of two replicates. Assays were performed in triplicate with similar results.

**Supplementary Figure 5.**
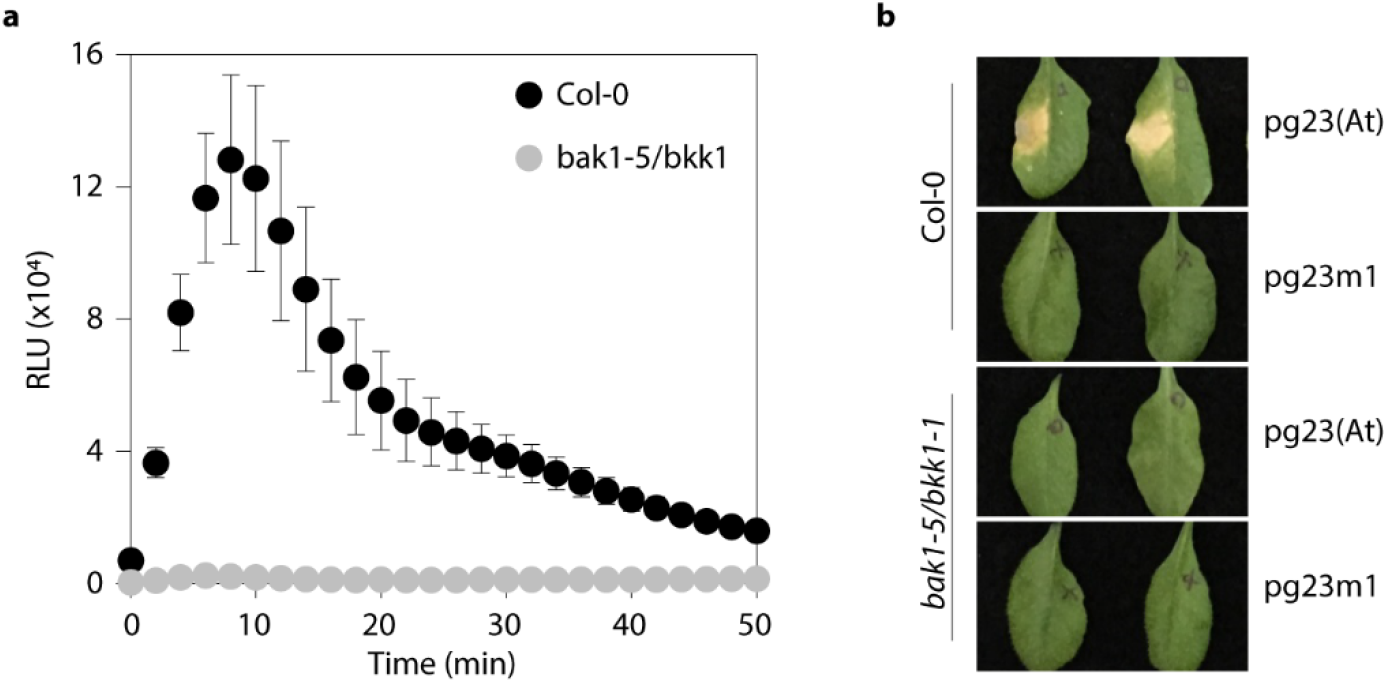
BAK1 and BKK1 are required for RLP42-pg9(At) signaling. **a**, ROS production (relative light units, RLU) in leaf discs of Arabidopsis Col-0 and *bak1-5/bkk1-1* mutant treated with 1 μM pg9(At). **b**, Hypersensitive-like cell death in leaves of Col-0 and bak1-5/bkk1-1 mutant infiltrated with 10 μM pg23(At) or pg23m1, and visualized at 7 days post infiltration.

**Supplementary Figure 6.**
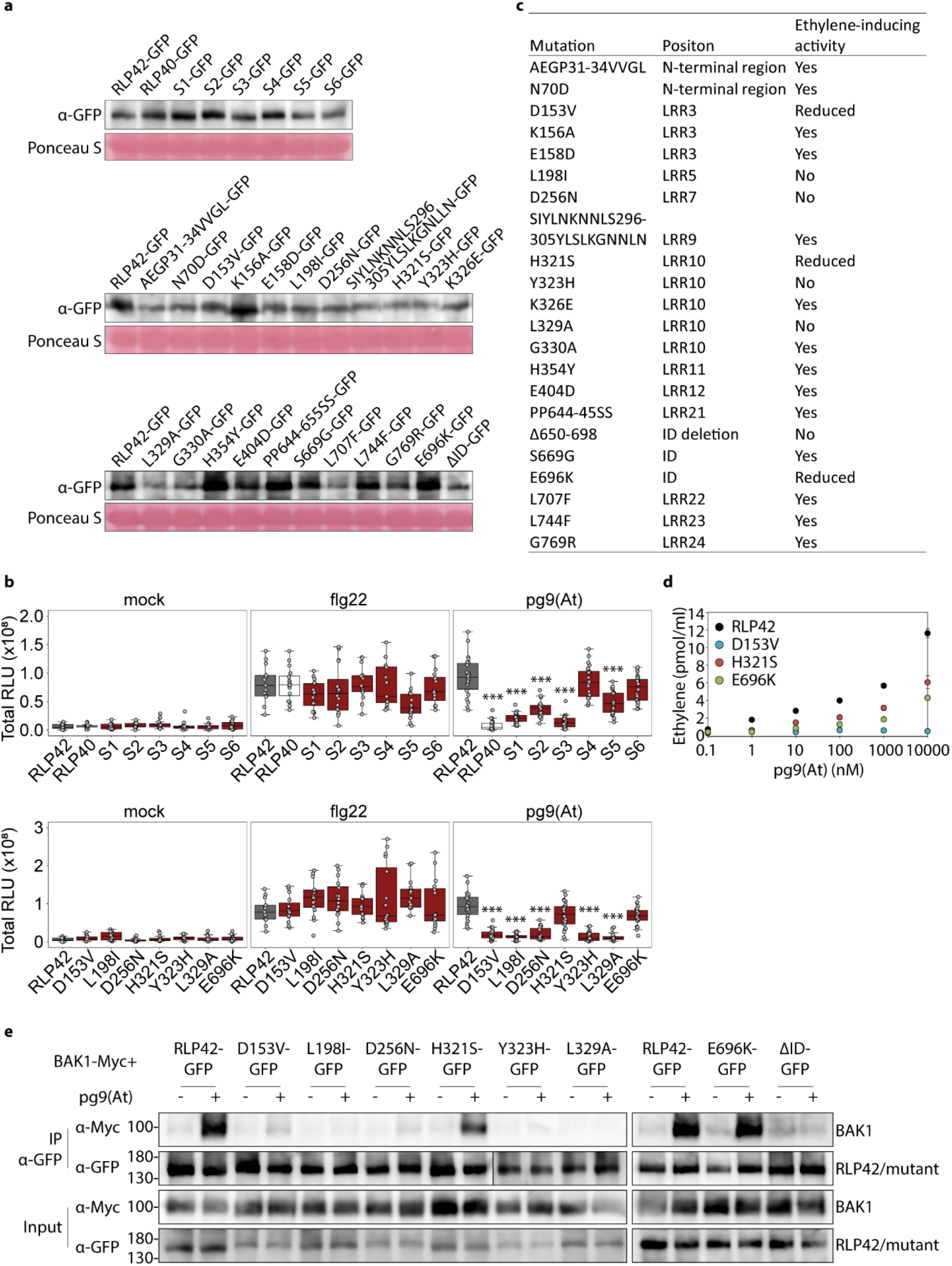
Structure-function analysis of RLP42 required for pg9(At) recognition. **a,** Immunoblots of transiently expressed RLP42 chimeric and mutant proteins in *Nicotiana benthamiana* leaves with anti-GFP antibody. **b,** Total ROS production (relative light units, RLU) in leaf discs of *N. benthamiana* transiently transformed with the chimeric constructs or point/short stretch mutation constructs with 1 μM pg9(At) or flg22, or water (mock). Data points are indicated as grey dots from three independent experiments (20≤n≤24) and plotted as box plots. (*** *P*<0.001, Student’s *t*-test). **c**, Summary of RLP42 chimeric and mutation proteins used in this study. **d,** Ethylene production after treatment with serial dilutions of pg9(At) in *N. benthamiana* leaves transiently transformed with RLP42, D153V, H321S, or E696K constructs. Bars represent means ± standard deviation of three replicates. Three independent experiments were performed with similar results. **e,** BAK1 recruitment to RLP42 receptor mutant proteins. Proteins extracted from *N. benthamiana* leaves co-expressing RLP42/mutant-GFP with BAK1-Myc, and treated with water (−) or 1 μM pg9(At) (+) for 5 min before harvesting, were used for co-immunoprecipitation with GFP-trap beads, and immunoblotting with tag-specific antibodies. Assays were performed in triplicate with similar results.

**Supplementary Figure 7.**
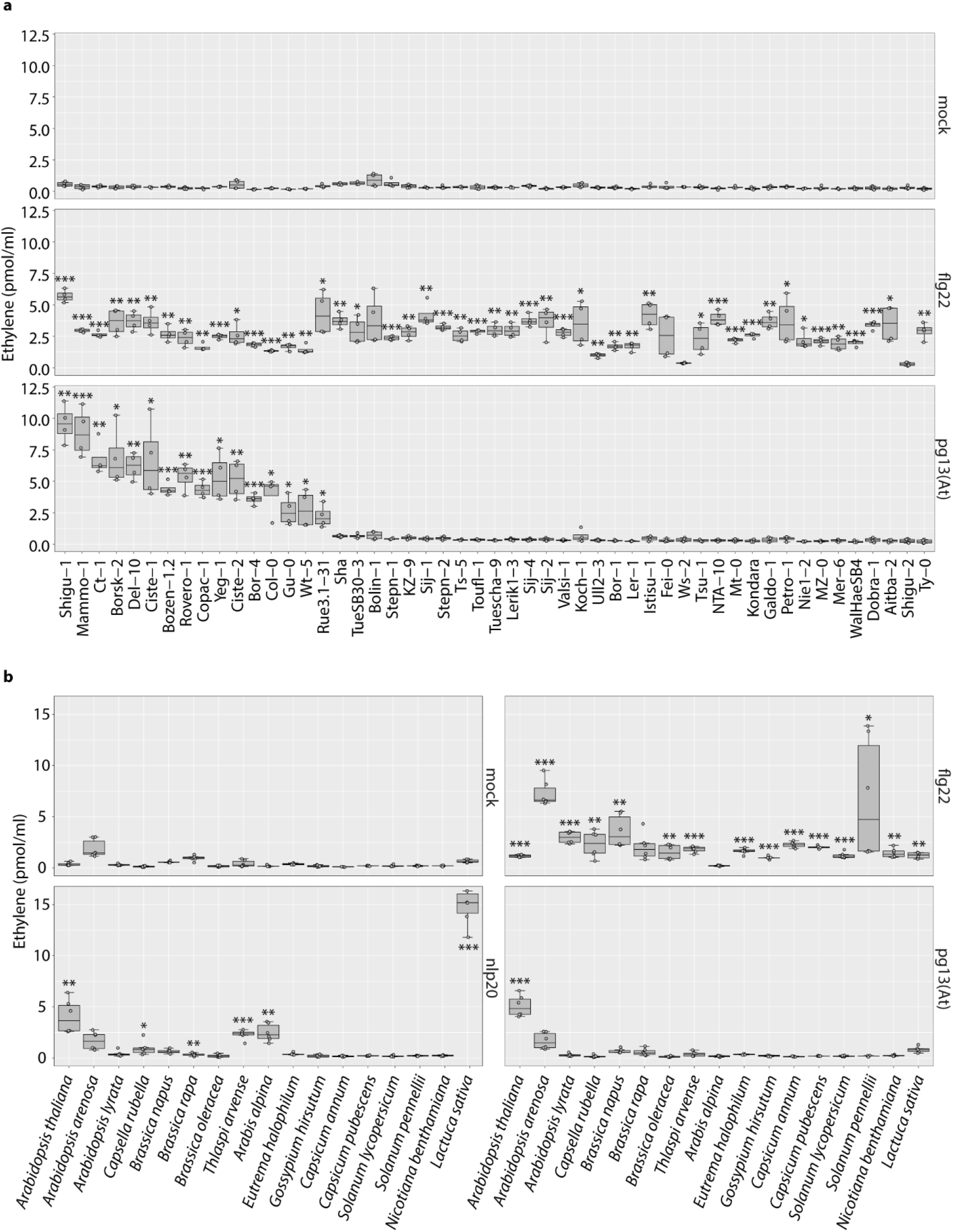
Pattern responsiveness in *Arabidopsis thaliana* accessions and different plant species. **a**, Ethylene accumulation in *A. thaliana* accessions after 4 h treatment with water (mock) or 1 μM flg22 or pg13(At). **b**, Ethylene accumulation in different plant species after 4 h treatment with water (mock) or 1 μM flg22, nlp20 or pg13(At). Data points are indicated as grey dots from two independent experiments (**a**, n=4; **b**, n=6) and plotted as box plots. Asterisks indicate statistical differences to mock treatments in respective plant (*** *P*<0.001, ** *P*<0.01, * *P*<0.05, Student’s *t*-test).

**Supplementary Figure 8.**
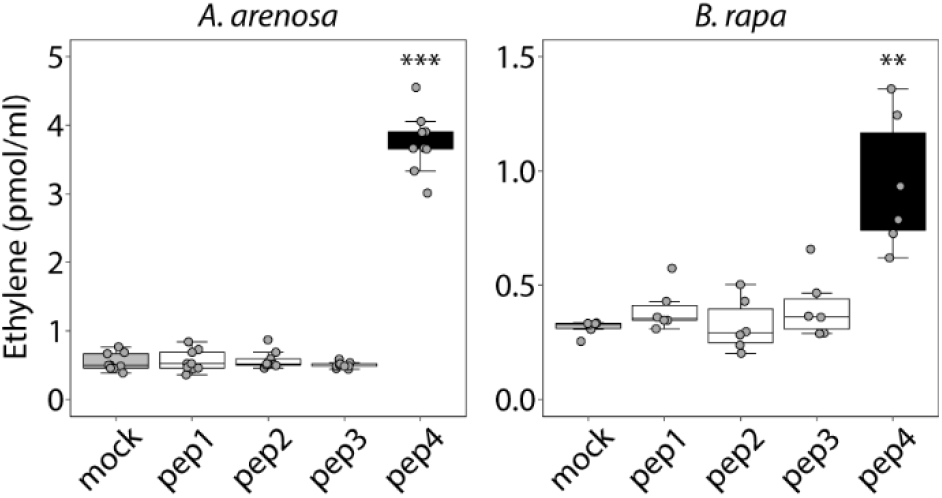
Pep1-3 derived from PG6 do not activate plant immune responses in *Arabidopsis arenosa* and *Brassica rapa*. Ethylene accumulation after 4 h treatment with water (mock), or 1 μM pep1-3 in *A. arenosa* and *B. rapa*. Data points are indicated as grey dots from three or two independent experiments (*A. arenosa*, n=9; *B. rapa*, n=6) and plotted as box plots. Asterisks indicate significant differences to mock treatments in the respective plant (*** *P*<0.001, ** *P*<0.01, Student’s *t*-test).

**Supplementary Figure 9.**
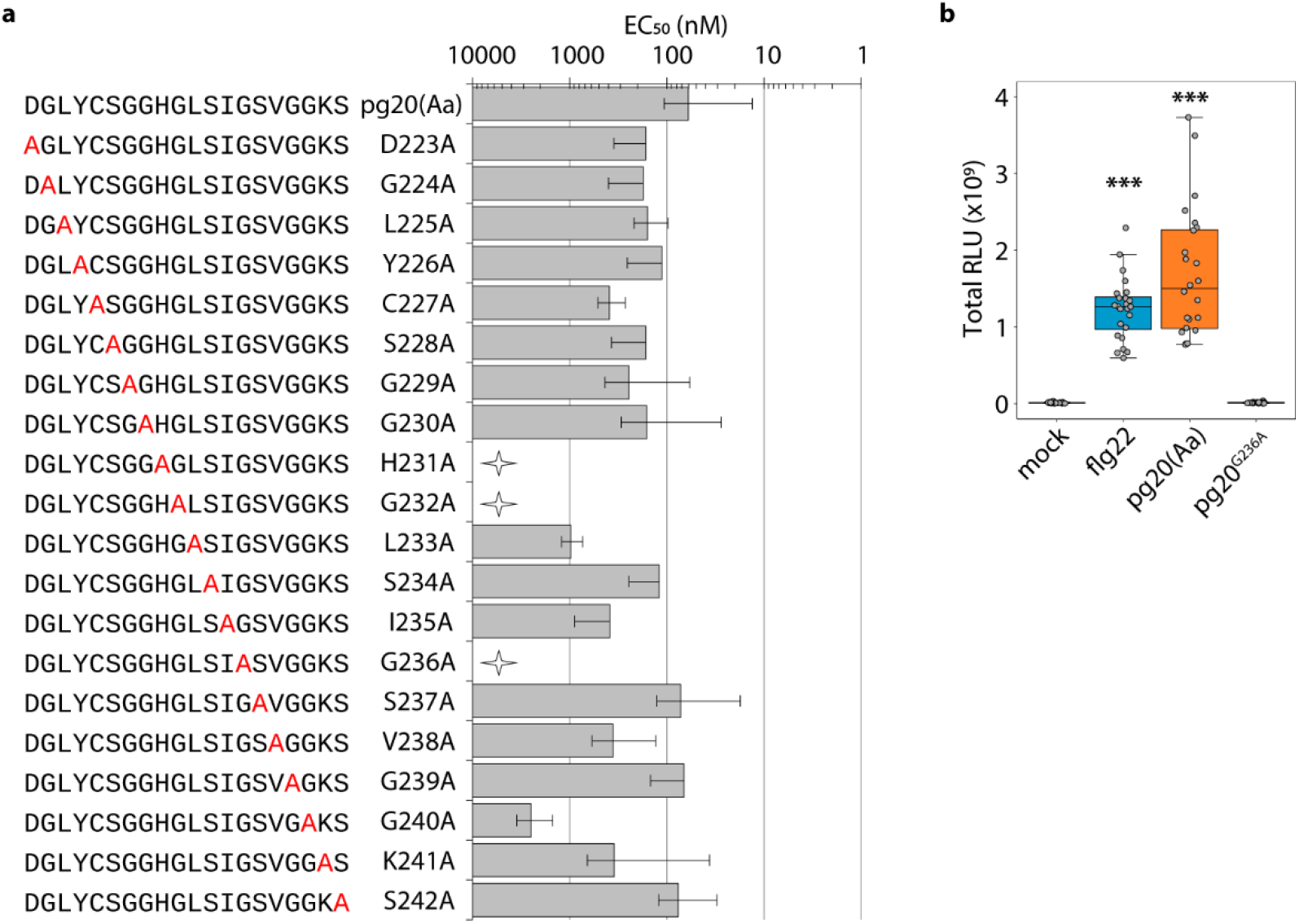
Pg20(Aa) activates plant immune responses in *Arabidopsis arenosa*. **a**, Ethylene-inducing activity of pg20(Aa) and the corresponding mutant peptides in *A. arenosa*. EC_50_ values were determined from dose-response curves with the synthetic peptides. Peptide sequences are indicated at the left panel. The mutant residues are indicated in red. At the right panel, bars represent means ± standard deviation on a logarithmic scale from at least three independent biological repeats, each with three replicates. The peptides that did not induce any or residual ethylene production only at 10 μM are defined as inactive peptide (stars). **b**, Total ROS production in leaf discs of *A. arenosa* treated with water (mock), 100 nM flg22, or 1 μM pg20 or pg20^G236A^ over 120 min. RLU, relative light unit. Data points are indicated as grey dots from three independent experiments (n=24) and plotted as box plots. Asterisks indicate significant differences to mock treatments (*** *P*<0.001, Student’s *t*-test).

**Supplementary Figure 10.**
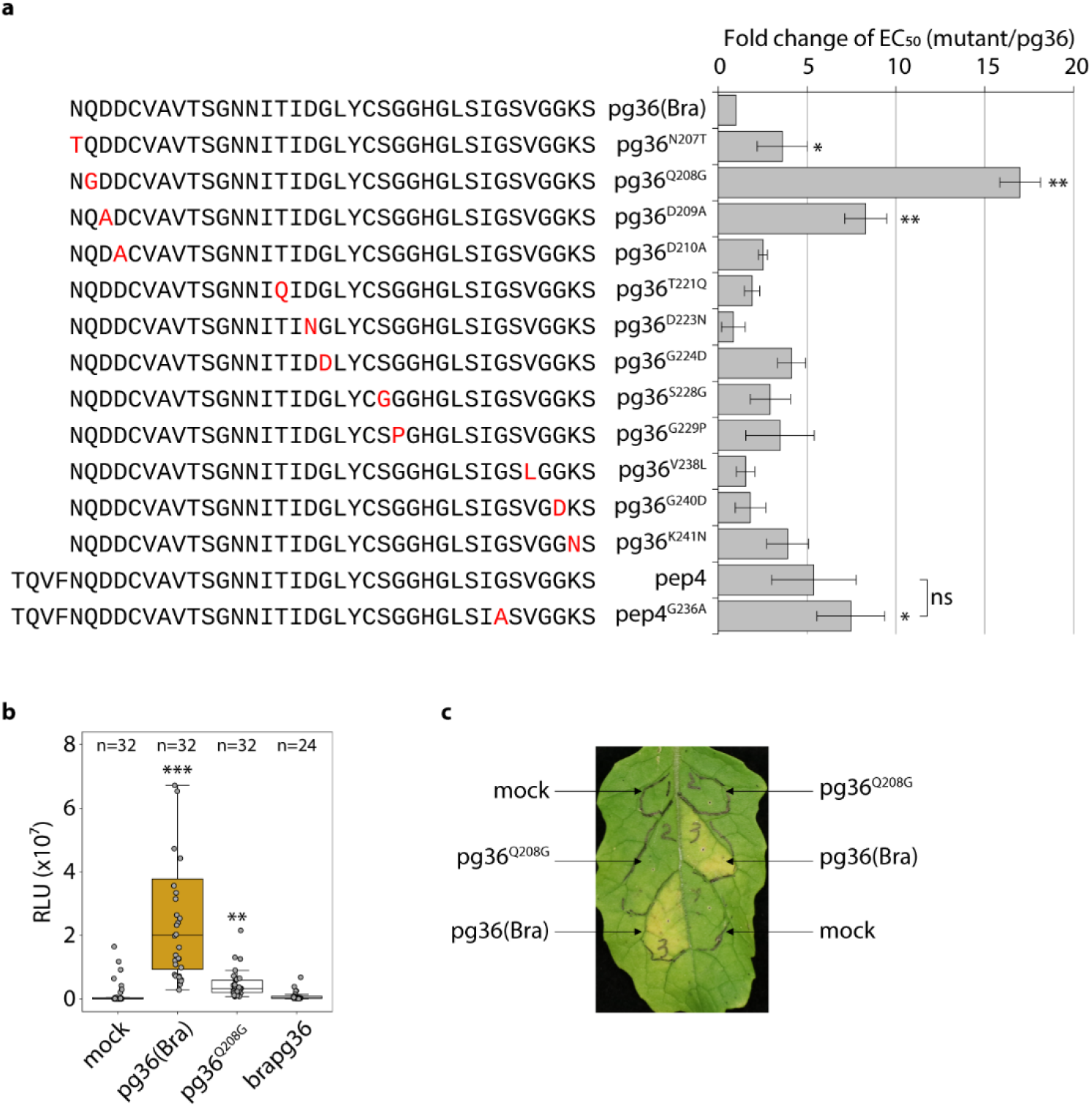
Pg36(Bra) activates plant immune responses in *Brassica rapa*. **a**, Ethylene-inducing activity of pg36(Bra) and the corresponding mutant peptides in *A. arenosa*. EC_50_ values were determined from dose-response curves with the synthetic peptides. Fold change of mutant peptide EC_50_ to that of pg36(Bra) was calculated. Peptide sequences are indicated at the left panel. The mutant residues are indicated in red. At the right panel, bars represent means ± standard deviation from at least three independent biological repeats, each with three replicates. **b**, Total ROS production in leaf discs of *B. rapa* treated with water (mock), or 1 μM of the given elicitor over 120 min. RLU, relative light unit. Data points are indicated as grey dots from at least three independent experiments and plotted as box plots. **c**, Hypersensitive-like cell death in leaves of *B. rapa* leaves infiltrated with 1 μM pg36(Bra) or pg36^Q208G^, or water (mock) and visualized at 7 days post infiltration. Assays were performed in triplicate with similar results. Asterisks indicate significant differences to pg36(Bra) (**a**) or mock (**b**) treatment ((*** *P*<0.001, ** *P*<0.01, * *P*<0.05, ‘ns’ no significant difference, Student’s *t*-test).

**Supplementary Table 1.**
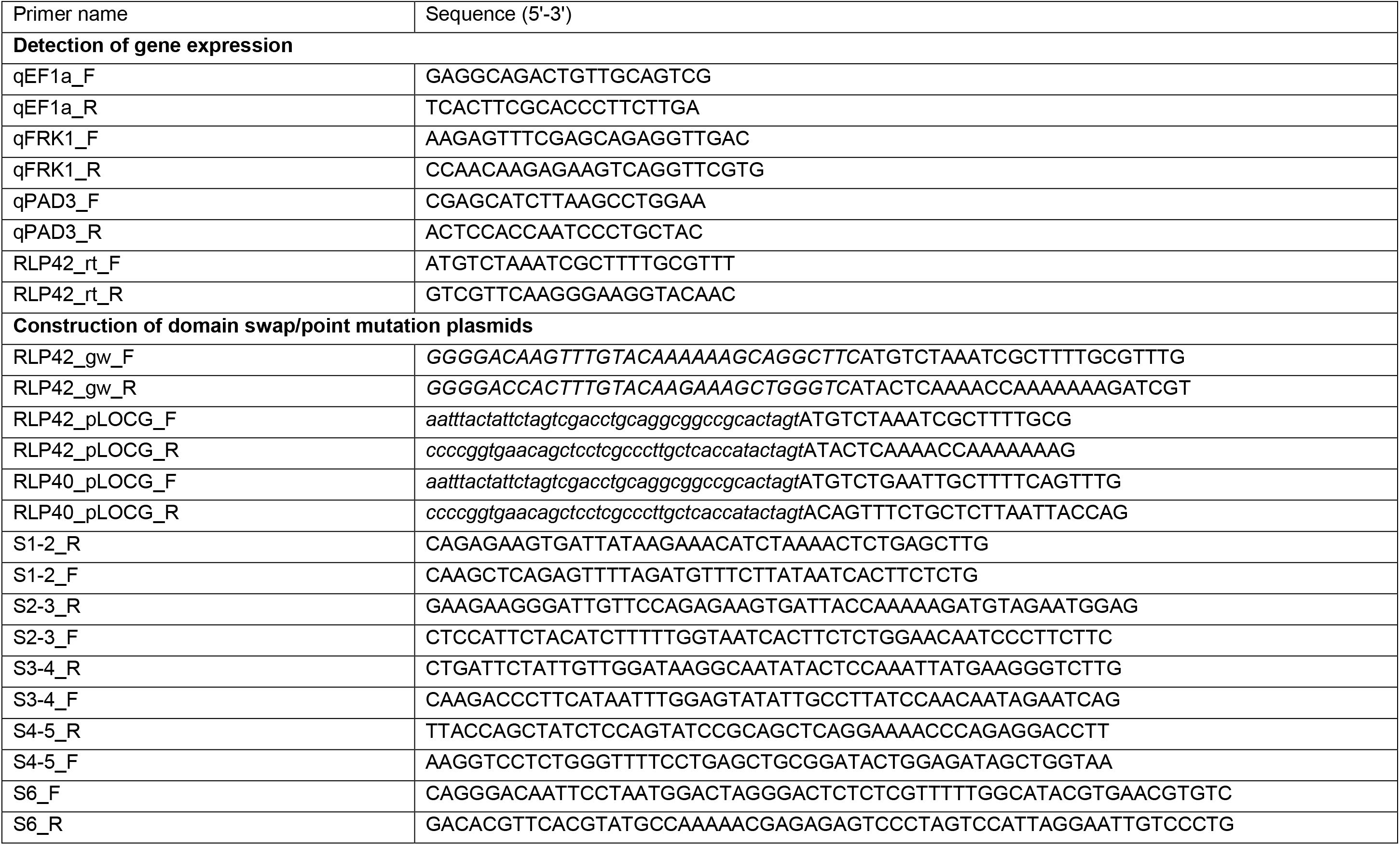
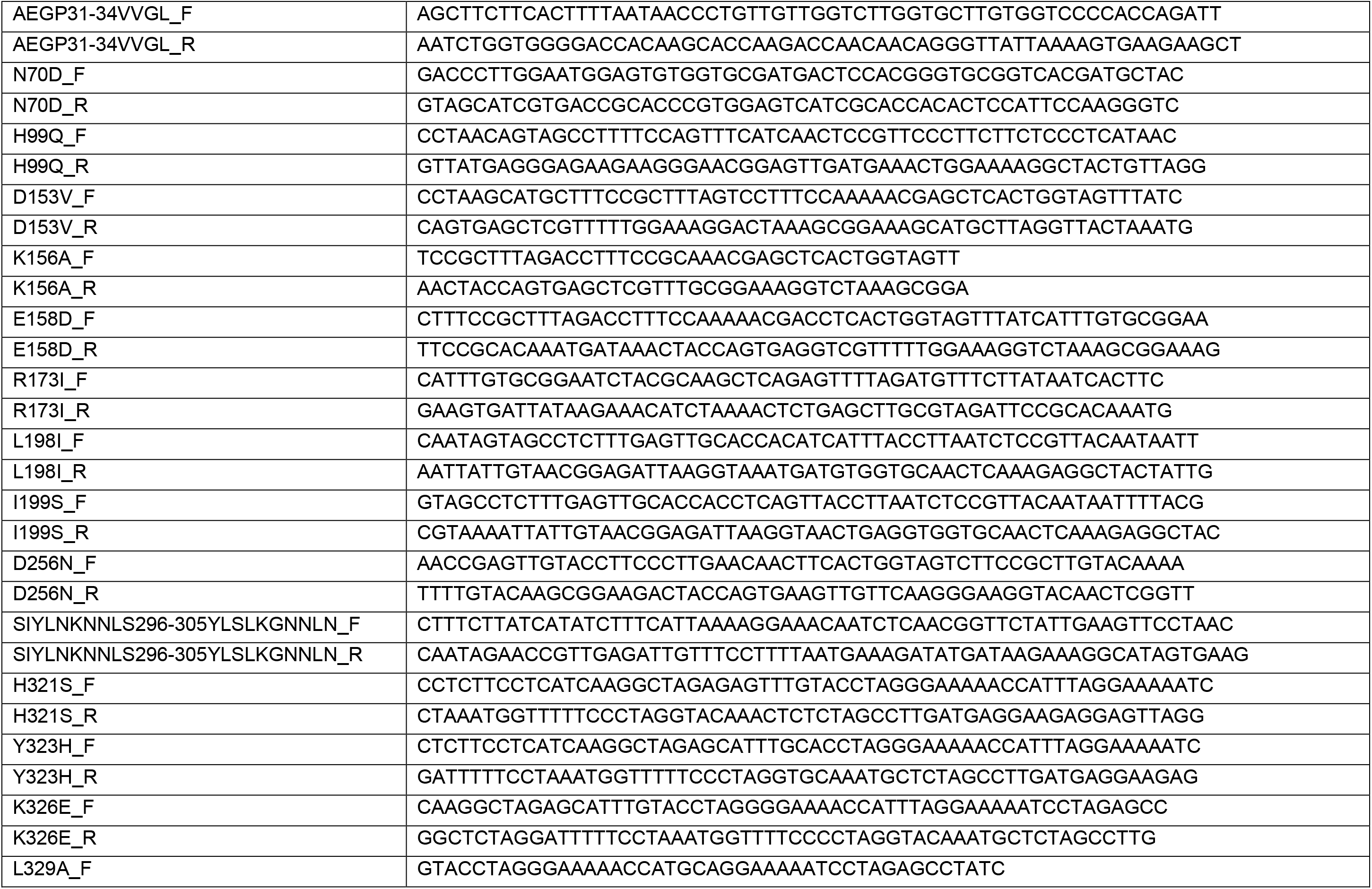
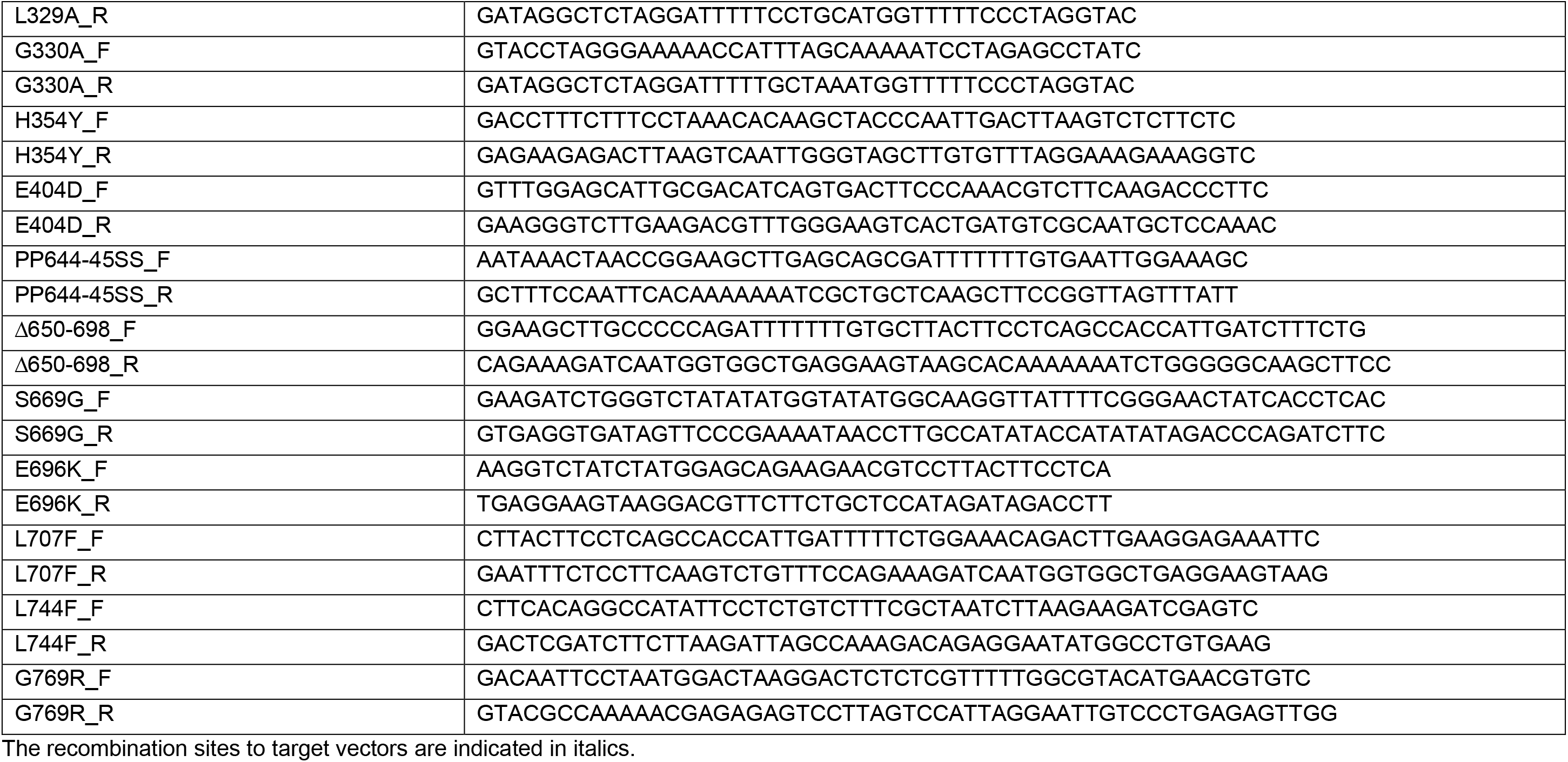
Primers used in this study.

